# Semantic search using protein large language models detects class II microcins in bacterial genomes

**DOI:** 10.1101/2023.11.15.567263

**Authors:** Anastasiya V. Kulikova, Jennifer K. Parker, Bryan W. Davies, Claus O. Wilke

**Affiliations:** Department of Integrative Biology, University of Texas at Austin, Austin, Texas, USA; Department of Molecular Biosciences, The University of Texas at Austin, Austin, TX, USA; John Ring LaMontagne Center for Infectious Diseases, The University of Texas at Austin, Austin, TX, USA

## Abstract

Class II microcins are antimicrobial peptides that have shown some potential as novel antibiotics. However, to date only ten class II microcins have been described, and discovery of novel microcins has been hampered by their short length and high sequence divergence. Here, we ask if we can use numerical embeddings generated by protein large language models to detect microcins in bacterial genome assemblies and whether this method can outperform sequence-based methods such as BLAST. We find that embeddings detect known class II microcins much more reliably than does BLAST and that any two microcins tend to have a small distance in embedding space even though they typically are highly diverged at the sequence level. In datasets of *Escherichia coli*, *Klebsiella* spp., and *Enterobacter* spp. genomes, we further find novel putative microcins that were previously missed by sequence-based search methods.

## 1 Introduction

The increasing problems of antibiotic resistance, failure of first-line treatments for bacterial infections, and a limited pipeline of drugs under development suggest that novel approaches to fighting bacterial infections may be required [1; 2]. One class of potential novel therapeutics are antibacterial proteins and peptides, which are found across all forms of life and are very diverse in structure and function [3; 4]. Among these are class II microcins, a class of antibacterial peptides produced by gram-negative bacteria which have shown some therapeutic potential *in vivo* [5; 6]. Microcins are part of a larger group of antibacterial proteins, called bacteriocins, which are produced by bacteria and toxic towards other bacteria. Mature class II microcins are less than 100 amino acids and 10 kDa, and their diversity and mechanisms of action are still largely unexplored [7]. The larger, proteinaceous bacteriocins have recently become more popular as potential antibacterial therapeutics, in part due to rapid progress in protein sequencing and the expansion of bioinformatics data [8; 9; 10]. Microcins also have potential to be used as antibiotics, especially as more research on their structure and function becomes available.

Unfortunately, only ten class II microcins with partial structural and/or functional characterization have been published to date. An early effort to screen for gram-negative bacterial peptides containing “double-glycine” signal sequences *in silico* may have identified microcins but was not explicitly designed to do so [11]. More recently, our group designed a computational pipeline specifically for the detection of class II micocins [12]. This pipeline, cinful, relies on both BLAST and a profile hidden Markov model (pHMM) trained on the ten published class II microcin sequences [12]. The application of cinful to over a thousand Enterobacterales genomes successfully expanded the sequence range of putative microcins. However, because of the limited number of experimentally verified microcins and the high sequence diversity among them, cinful may not work optimally for detecting the full range of novel, previously unidentified microcins that exist. Moreover, since microcins have high sequence divergence, there is a need for alternative approaches that do not primarily rely on alignments to detect microcins.

Here, we investigate an alternative method of identifying putative microcin sequences. Instead of employing sequence similarity as the primary search tool, we are instead searching for open reading frames (ORFs) that are close to known microcins in a high-dimensional embedding space generated by the protein large language model, ESM-1b [13; 14]. First, we show that microcins cluster together in embedding space when compared to non-microcin ORFs. Second, we show that almost every single one of the ten known microcins can be used to recover all other known microcins from their genomic background using distance in embedding space. By contrast, BLAST searches perform poorly at the same task. Third, we use embeddings to search through *Escherichia coli*, *Klebsiella* spp., and *Enterobacter* spp. genomes where prior, sequence-based methods found a microcin exporter gene but no actual microcins. In these genomes, our approach identifies novel putative microcin sequences.

## 2 Methods

### 2.1 Data collection

We collected seven genomes, plasmids, or gene clusters (Table 1) that contained the ten known microcins. These microcins are class IIa microcins V [15], L [16], N [17], PDI [18], and S [19]; and class IIb microcins H47 [20], I47 [21], M [22], E492 [23], and G492 [24]. The genomic sequences containing these microcins were used to validate embedding distance as a tool to search for microcin sequences in genomic data.

**Table 1:** Genomes, gene clusters, or plasmids containing the ten known microcins. The column “extracted ORFs” refers to the number of putative open reading frames remaining after filtering the initial extracted ORFs by size.

| genome/cluster | accession | extracted ORFs | microcins |
| --- | --- | --- | --- |
| *E. coli* pColV-K30 | AJ223631.1 | 157 | V |
| *E. coli* NCTC11128 | GCF_900448705.1 | 25819 | N |
| *E. coli* 54909 | GCF_025261725.1 | 27746 | L |
| *K. pneumoniae* cluster | AF063590.3 | 372 | E492, G492 |
| *E. coli* H47 cluster | AJ009631.3 | 196 | I47, H47, M |
| *E. coli* G3/10 plasmid | JN887338.1 | 258 | S |
| *E. coli* 25 plasmid | JQ901381.1 | 647 | PDI |

To search for novel class II microcins, we compiled three datasets containing 25 *E. coli*, 44 *Enterobacter* spp., and 46 *Klebsiella* spp. genomes, respectively (Supplementary Files S1, S2 and S3). When choosing the genomes for each dataset, we looked at genomes where the previous method, cinful, did not find any microcins, yet found the peptidase-containing ABC transporter (PCAT) microcin exporter gene [12]. These genome datasets were taken from a subset of the GTBD genomes that were used in [12]. Cinful results can be found in Supplementary File S4). We next extracted ORFs from each genome, generated embeddings for each ORF, and searched each of the 25 genomes using the embeddings from each of the ten known microcin sequences.

We also searched through 40 *E. coli* genomes from the B2 phylogroup from [25] (Supplementary File S5). Of these 40 genomes, 20 were from *E. coli* isolated from water samples, 15 of which had no previously detected microcins [12]. The other five were the only genomes from water isolates where microcins had been found by cinful [12]. The other 20 genomes were from *E. coli* isolated from human extra-intestinal samples; we selected 10 genomes where putative microcins had been found and 10 genomes where no microcins had previously been found [12].

### 2.2 Extracting ORFs and generating embeddings

We used a simple method to identify putative ORFs in each genome or gene assembly: we identified all contiguous stretches of codons starting with ATG (methionine) and ending with one of the three stop codons. We further imposed a minimum size limit of 30 residues and a maximum size limit of 150 residues, since all known microcins fall within these limits. In cases where ORFs had multiple possible start codons, we we only kept the longest ORF within the size limit stated above. This process will recover many putative ORFs that do not correspond to known protein-coding genes. We accepted this high false-positive rate in our ORF-finding procedure since our goal was to detect as many putative microcins as possible and some of them might be missed under a more restrictive definition of an ORF. For each genome, we searched for ORFs in all possible reading frames starting from methionine, both on the + and the − strand.

We generated ESM1-b embeddings for each ORF using the ESM library available at: https://github.com/facebookresearch/ESM. Embeddings were initially generated for each residue in a sequence. Each embedding consists of a numeric vector of 1280 features. To arrive at a single embedding per ORF, we subsequently averaged each embedding feature (one of the 1280 numbers) across all residues in the sequence. The resulting averaged embedding is a numeric vector of 1280 values representing the entire ORF. We generated these averaged embeddings for all extracted ORFs and all known microcins.

### 2.3 Principal component analysis

For the principal component analysis (PCA), we used 27,745 putative ORFs of microcin length (30-150 amino acids) from the *E. coli* 54909 genome. The ORFs were extracted and converted into embeddings as described above. The *E. coli* 54909 genome contains the known microcin L, which we removed from the putative ORFs to arrive at the non-microcin ORF group. For the microcin group, we used the embeddings for the ten known microcins.

### 2.4 Calculating distance and identifying microcin hits

The mean protein embeddings are vectors of *n* = 1280 features. The distance between two embeddings was calculated as the Euclidean distance between two points. Specifically, assuming two embedding vectors *p* and *q*, we calculated the distance 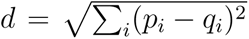, where *p_i_* and *q_i_* represent the *i*th component of each of the two embeddings.

After calculating the distances between a query microcin embedding and the embedding for each ORF in the target genome, we plotted ORFs in order of ascending embedding distance and manually inspected the lowest 50 distances. For each target genome, we repeated this process 11 times, once for each of the 10 known microcins and once for the average embedding of all 10 microcins. Most genomes will generally contain one to two microcins, and it is rare for an assembly to have more that two microcins [12]. For this reason, we inspected the two ORFs with the lowest embedding distance to any of the queries in each genome. If any one or two ORFs displayed a meaningful gap from the remainder of the distribution of distances and if they were found to be one of the two lowest distance ORFs by more than one query microcin, these one/two ORFs were declared “hits” and further inspected for microcin features.

Multiple sequence alignments of microcin sequences were performed using MAFFT from the EMBL-EBI search and sequence analysis tools [26], availabe at: https://www.ebi.ac.uk/Tools/msa/mafft/. Alignments were visualized using MView from the EMBL-EBI search and sequence analysis tools, available at: https://www.ebi.ac.uk/Tools/msa/mview/.

### 2.5 Running BLAST

To run BLAST searches, we first took all extracted ORFs from the respective genome or gene assembly to be searched and converted them into a BLAST database. We then used each of the 10 known microcin sequences as a query against each of these databases.

All queries were run with protein BLAST v2.13.0 using an E-value cut-off of 10 and a maximum of 50 target sequences. After generating initial BLAST results, we further filtered the results to only keep hits where the hit alignment length was *≥* 30 residues and the percent mismatch was *<* 80%. We then checked these remaining ORF hits to see if they were or contained known microcin sequences.

### 2.6 Calculating percent sequence divergence

Percent sequence divergence was calculated by first aligning a pair of sequences and then counting the total number of sites that did not match and dividing by the length of the alignment. Any gaps in the alignment were also treated as mismatches. All pairwise alignments were performed using gemmi (v0.5.7) [27].

To analyze sequence divergence and embedding distance of *E. coli* ORFs, we used the ORFs extracted from the *E. coli* 54909 genome containing microcin L. We first calculated sequence divergence for 112,017,666 arbitrarily selected non-microcin ORF pairs, which corresponds to approximately 15% of all total possible pairs. The initial sample was chosen by systematically calculating sequence divergence for all possible ORF pairs until we had collected a large number of pairs spanning a wide range of sequence divergence. We then further down-sampled this dataset for visualization purposes. The goal of our down-sampling procedure was to obtain uniform representation of the entire range of possible sequence divergence values. To this end, we binned all data into bins spanning 10 percentage points in sequence divergence. Because low-divergence ORF pairs are rare, we kept all data within the lowest three divergence bins from 0–30% divergence. Next, since the 30– 40% divergence group only had 25 pairs, we kept those pairs and arbitrarily selected 25 pairs for the remaining higher divergence bins. In the final sample, there were a total of 194 pairs; 175 pairs between 100 and 30% divergence (25 per 10% bin), nine pairs with divergence between 20–30%, four pairs between 10–20% and six pairs between 0 and 10%. After down-sampling, we generated embeddings for all ORFs in the pared dataset and calculated the embedding distance for all ORF pairs. We also calculated sequence divergence and embedding distance for all possible combinations of the 10 known microcins, resulting in data for 45 pairs.

### 2.7 Data availability

All data analysis and figure production were performed in R [28], using the tidyverse family of packages [29]. Trained neural networks, analysis scripts, and processed data are available on GitHub: https://github.com/akulikova64/microcin_embeddings_project

## 3 Results

We wanted to know whether semantic embeddings from a protein large language model (LLM) can be used to search for small proteins in bacterial genomes. Specifically, we were interested in detecting class II microcins, which are short (fewer than 150 amino acids) and highly diverged, making them difficult to find using traditional sequence-based approaches.

To assess the overall feasibility of our approach, we first asked whether embeddings for microcins cluster together when compared to embeddings for non-microcin ORFs. We generated embeddings for the 10 currently known microcins (Supplementary File S6), using the protein large language model, ESM1-b [13]. ESM1-b is a widely-used LLM trained on 250 million known protein sequences. The complete set of ESM1-b embeddings for a single protein sequence of length *L* is a matrix of 1280 rows and *L* columns. To reduce this data to a more manageable set of numbers, for each protein we calculated the average across all *L* columns for each row and used the resulting set of 1280 averages as our “semantic embedding” vector representing various characteristics of the biochemistry of the input protein.

For comparison, we similarly calculated semantic embedding vectors for all putative non-microcin ORFs of length 30–150 amino acids in the *E. coli* 54909 genome, for a total of 27,745 ORFs. We chose this genome as it is one of the two complete *E. coli* genomes in which microcin L is found. The large number of putative ORFs is due to our ORF extractor, which finds all contiguous stretches of non-stop codons beginning with ATG and within the correct size range (see Methods). This definition includes many low-confidence ORFs. For comparison, a common *E. coli* K-12 genome contains about 4,333 high-confidence ORFs. We then performed a principal component analysis (PCA) on the combined set of embeddings (microcin and non-microcin ORFs) and plotted the location of each ORF within the first two components (Figure 1). We found that all microcins were tightly clustered within a small region of the overall embedding space, suggesting that microcins share similar characteristics that the embeddings can capture. However, non-microcin ORFs were similarly present in the same region of principal component space, i.e., the microcins did not trivially separate from the remainder of the ORFs in the genome. We emphasize that the full embedding space has 1280 dimensions, so a more sophisticated approach considering all dimensions can potentially separate microcin ORFs from non-microcin ORFs even if they overlap in the two-dimensional PCA plot. More importantly, in the PCA plot the microcin embeddings fell along the periphery of the ORF embedding distribution and did not overlap with the majority of non-microcin embeddings.

**Figure 1:**
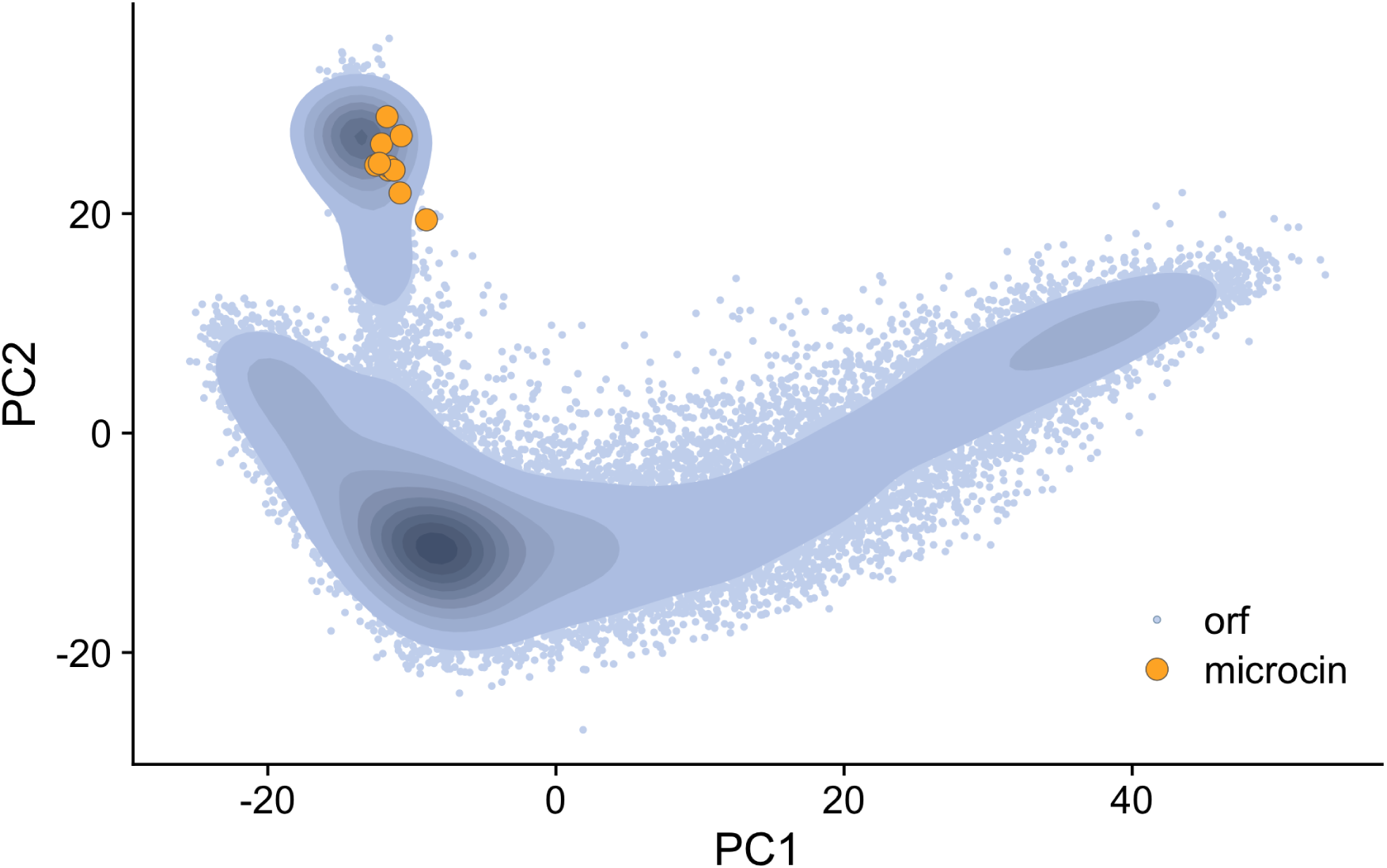
PCA plot of *E. coli* 54909 non-microcin ORFs and the 10 known microcins. Blue points represent non-microcin ORF embeddings, while the orange points represent embeddings of the 10 known microcins. Darker colors along the blue gradient show higher density of points for non-microcin ORFs. There are a total of 27745 non-microcin ORFs.

Next, we tested whether or not each of the 10 known microcins can be used in a database search to find all other known microcins among a collection of ORFs extracted from either a genome or a gene cluster. We collected assemblies of genomes, gene clusters, or plasmids, as appropriate, which contain one or more of the known microcin sequences (Table 1). We extracted all ORFs for each of these assemblies (Figure 2a). Because the known microcins range from 75–120 amino acid residues long, we discarded all ORFs that were either longer than 150 residues or shorter than 30 residues. We then generated ESM1-b embeddings for all extracted ORFs (Figure 2c). Each of the 10 microcin embeddings were consecutively used as the query sequence to search within each assembly (Figure 2b). As an additional query, we used the average embedding of the 10 microcin embeddings. We calculated the distance between each ORF embedding in the assembly to the query microcin embedding and generated a set of distances for each query embedding (Figure 2d). Finally, we plotted and ordered these distances within each set by ascending distance and only looked at the 50 smallest distances to the query (Figure 2e). In total, each assembly produced 11 distance plots; 10 for the query microcins and one for the averaged microcin embedding. We expected that potential microcin ORFs would have the smallest distance to the query embeddings and appear separated from the rest of the non-microcin ORF embeddings (orange points in Figure 2e and Figure 3)

**Figure 2:**
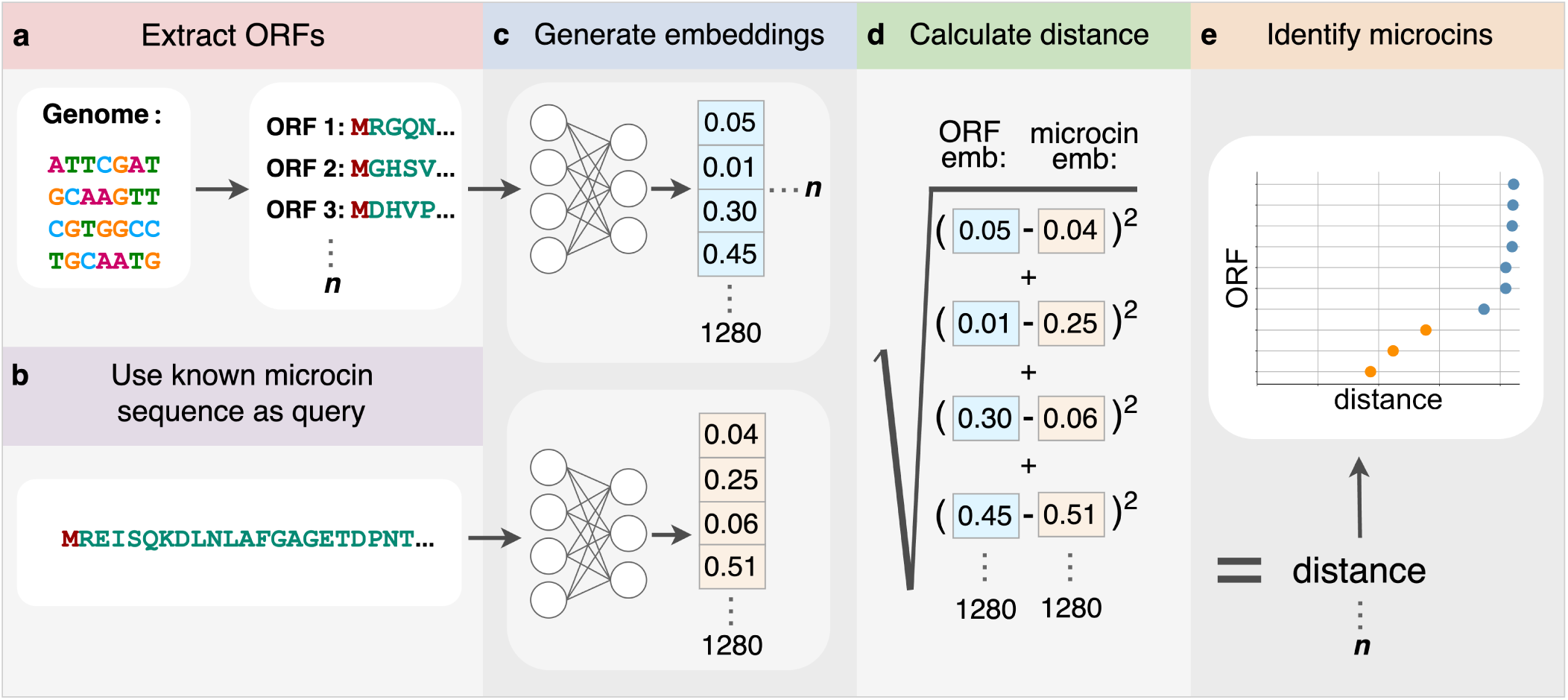
Detecting microcins using transformer embeddings. (a) Open reading frames (ORFs) are extracted from either whole genomes, gene clusters, or plasmids. (b) Each microcin sequence is consecutively selected as the query. (c) Embeddings are generated for both the exctracted ORFs as well as the microcin query sequence. (d) Semantic distance is calculated as the Euclidean distance between the two embedding vectors. (e) Microcin ORFs (orange) are identified as a cluster of 1–3 points all having very small distances to the query microcin embedding.

**Figure 3:**
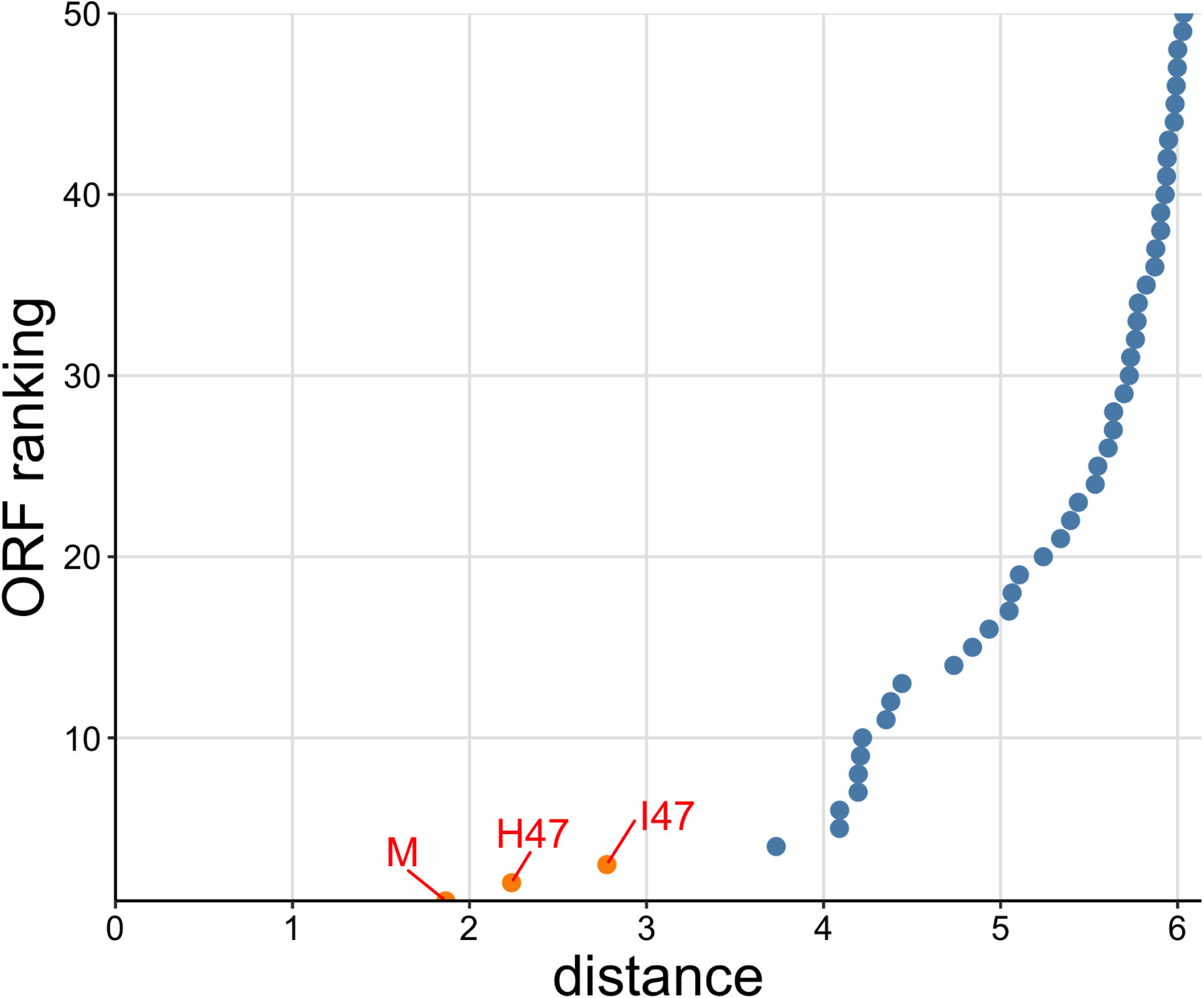
Search for microcins using microcin E492 as the query against the *E. coli* microcin H47 gene cluster. The microcin H47 gene cluster contains 405 extracted ORFs. Orange points indicate detected known microcins. Blue points indicate non-microcin ORFs. ORFs have been ranked based on increasing semantic distance.

To analyze the performance of this search algorithm more systematically, we defined a microcin as “found” if its distance was within the five lowest distances to the query (orange in Figure 2e), and “not found” otherwise. We found that nearly every microcin query sequence was able to recover all of the 10 known microcins using this procedure (Figure 4a). Furthermore, in most cases, the microcin ORFs were in the top two hits, even though we considered the top five in our procedure. The main exception was microcin I47 when used as a query, which did not recover microcins G492, N, or L. Also, microcin G492 when used as a query did not recover microcin I47. We note that microcins I47 and G492 had the largest distance in embedding space among all possible microcin pairings (Supplementary Figure S1). We also emphasize that the distance in embedding space between two microcins is not by itself the sole determining factor in whether a given query recovers a given target. Microcins I47 and H47 had the second-largest distance in embedding space (Supplementary Figure S1), and either one could be used to recover the other (Figure 4a). Whether a known microcin is recovered within the five lowest distances depends not only on the distance from the query to that microcin but also on whether there are any other ORFs that happen to have a low distance in embedding space, regardless of whether or not they are potential microcins. In fact, in general it was more challenging to identify microcins (such as N and L) located in complete genomes with thousands of extracted ORFs rather than in small gene clusters with only hundreds of ORFs (see Table 1 for exact ORF numbers used in this study). Yet, all microcins other than I47 when used as the query were able to recover both microcins N and L.

**Figure 4:**
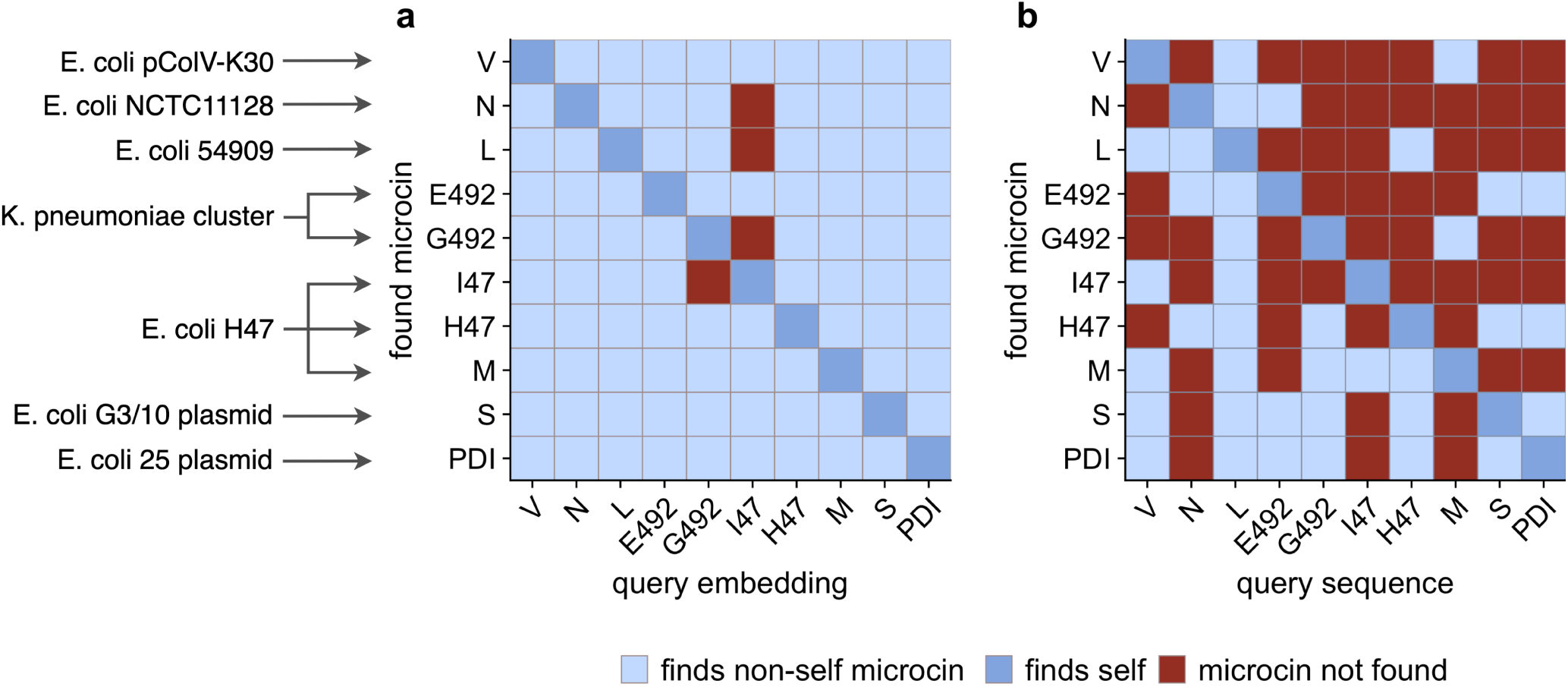
Performance of searches via semantic embeddings and via BLAST for known microcins across seven genomic assemblies. Light blue indicates that a microcin was found using a different microcin as the query. Dark blue indicates that a microcin was found using itself as the query. Red indicates that the microcin was not found. (a) When searching via semantic embeddings, nearly every individual known microcin can be found using any other known microcin as the query. (b) When searching via BLAST, most of the known microcins cannot be found using any other known microcins as the query. The only exception is the L microcin, which recovers all known microcins. BLAST parameters were as follows: E-value cutoff of 10 (default), alignment length *≥* 30 residues, percent mismatch *<* 80%.

We next contrasted our embedding-based search to a standard BLAST search. We took the same extracted ORFs we had used for the embedding method and employed them as the database sequences for BLAST. We then ran BLAST searches with each of the 10 microcin sequences as a query. We filtered all BLAST hits to retain only those hits with an E-value *≤* 10 and where the alignment length was at least 30 residues and the percent mismatch was less than 80%, rather non-restrictive parameters. Using this approach, very few microcin query sequences were able to recover the majority of the other microcin sequences. In fact, most microcins only recovered 2–3 other microcins. The one exception was microcin L, which was able to recover all other microcins (Figure 4b). It is unclear why microcin L performs so much better than the other microcins. In terms of sequence divergence, it shows similar values to any other microcin pairs. Nearly all known microcins are between 50%–70% diverged from any other microcin (Supplementary Figure S2).

BLAST has difficulty finding microcins because they are both short and highly diverged from each other. By contrast, distances in embedding space are small even for highly diverged microcins. To further explore the relationship between sequence divergence and distance in embedding space, we compared these distances for randomly chosen ORF pairs from the *E. coli* 54909 genome. Specifically, we picked 194 pairs of non-microcin ORFs (sampled at random such that they covered the entire range of sequence divergence values) and compared them to all possible microcin–microcin pairs. We found that, compared to non-microcin ORF pairs, microcins tended to have low semantic distance between each other given their sequence divergence, which was around 70% on average (Figure 5). Thus, in other words, even though microcins are highly diverged, they possess unique characteristics that can be picked up by the embedding space of a large language model and they can be uniquely identified by these characteristics.

**Figure 5:**
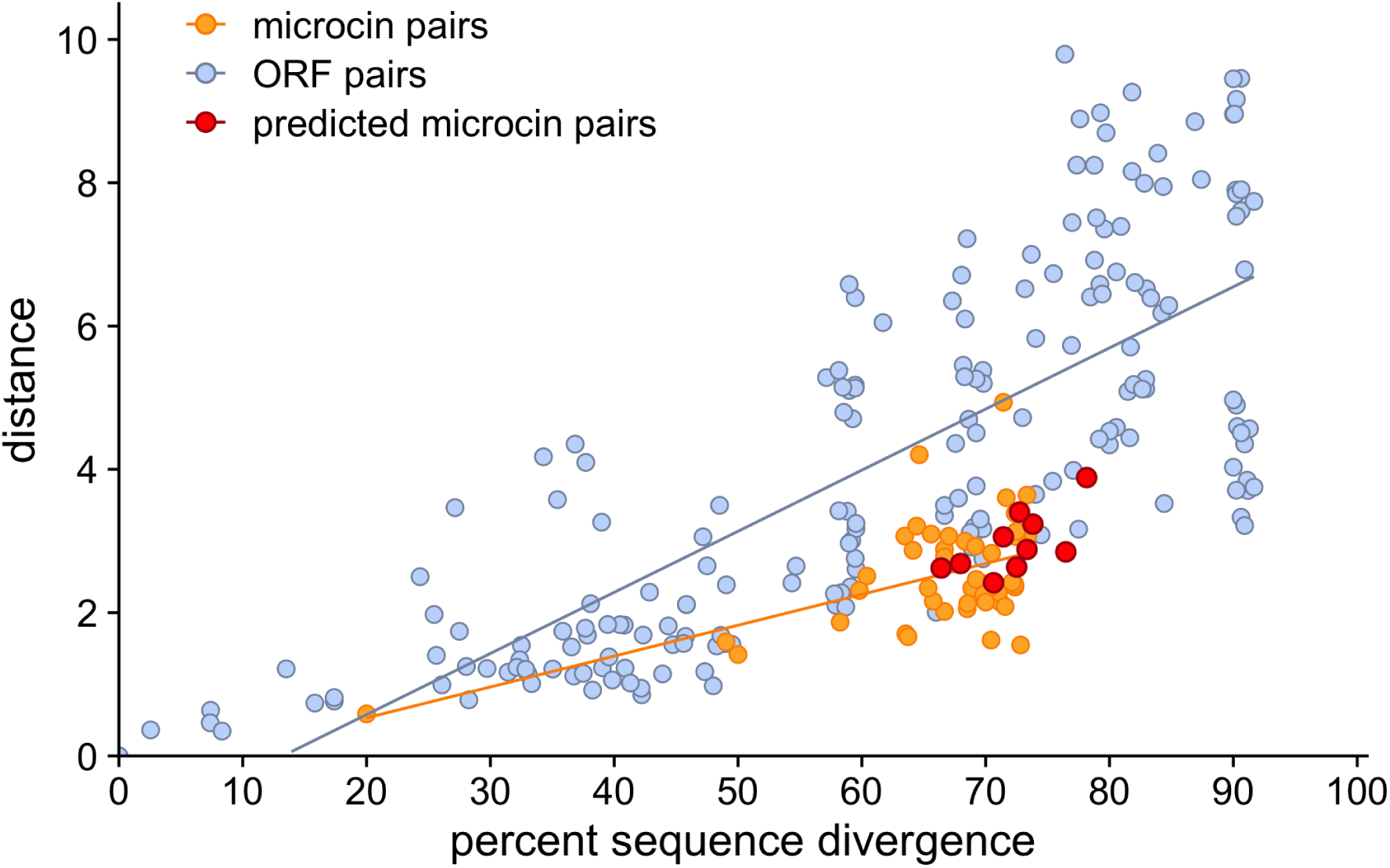
Semantic distance and percent sequence divergence for microcin pairs and non-microcin ORFs of comparable length. Blue dots indicate non-microcin ORF pairs, while orange dots indicate pairs of known pairs. For non-microcin ORF pairs, there are 194 pairs sampled across the range of sequence similarities. The microcin pairs include all 45 pairings of each of the 10 known microcins with every other one. The predicted microcin pairs include 10 pairings between each of the 10 known microcins and the putative microcin from *E. coli* TA097 identified in this work (see also Fig. 6).

Having established that embeddings may outperform sequence-based methods to locate microcins, we next asked whether we could discover novel putative microcins. In a previous study [12], we had used “cinful”, a sequence-based method for microcin discovery, to search for putative microcins in the Touchon dataset of *∼* 1000 *E. coli* genomes [25]. From this large dataset, we compiled a smaller dataset of 25 genomes where cinful had not previously found any putative microcins but did find the PCAT microcin exporter gene (Supplementary File S1). We reasoned that any genomes containing a microcin exporter would be prime candidates to contain microcins that may have been overlooked using the earlier methods.

For this systematic search for novel putative microcins, we proceeded as before, first calculating embeddings for all ORFs, then calculating distances between these embeddings and the embeddings of known microcins, and then looking for the ORFs with the smallest distances to known microcins. However, we slightly modified our definition of a hit to be more restrictive. We now only considered ORFs to be putative microcins if they were among the lowest two distances in the distribution of ORF distances for more that two query sequences and if there was also a meaningful gap between these distances and the rest of the distance distribution.

Furthermore, prior to analyzing any putative novel microcin hits, we needed to set criteria for what we would consider to be a likely true positive. Class II microcins have several characteristics that distinguish them from non-microcin peptides [7]. First, microcins contain a short (15–18 amino acid) “double-glycine” signal sequence at the N-terminus. This signal sequence terminates in either a glycine-glycine (GG) or glycine-alanine (GA) residue pair. Double-glycine signal sequences typically contain hydrophobic residues at positions −4, −7 and −12 relative to the signal-sequence cleavage site. These hydrophobic residues are critical for efficient secretion of double-glycine signal-containing peptides [30]. Class II microcin sequences are also unusually rich in glycine residues compared to other proteins, with glycine content ranging from 12–26%. Finally, class II microcins are encoded near their PCAT export gene on the genome. All of these characterisitcs were taken into account when identifying putative microcins among the output from the embeddings search.

Among the 25 genomes with a PCAT microcin exporter gene but no previously discovered putative microcins, our embeddings method detected a total of 44 hits (Supplementary File S7). Out of the 44 hits, three were identified as putative microcins (Table 2, Figure 6, and Supplementary File S8). One of those, microcin L located within the 36 1 Ti13 genome, had previously been missed because it had not been identified as an ORF by cinful’s ORF finder, Prodigal [31]. The more inclusive definition of ORFs we used here allowed us to find this microcin. The two other hits, however, were novel microcin-like sequences, discovered in the two genomes TA097 and TA265, respectively (Supplementary File S8 and Figure 6a). Both of these sequences displayed characteristics of microcins, such as high glycine content (yellow), a glycine-alanine (GA) residue pair before the cleavage site (pink) as well as hydrophobic residues (purple) at positions −4, −7 and −12 relative to the signal sequence cleavage site (Figure 6b). We also found that these two new putative microcins had similar sequence divergence and semantic distance to known microcins (Figure 5). Finally, we confirmed that these sequences had been identified as ORFs in the cinful pipeline and had been missed by the BLAST search employed by cinful.

**Figure 6:**
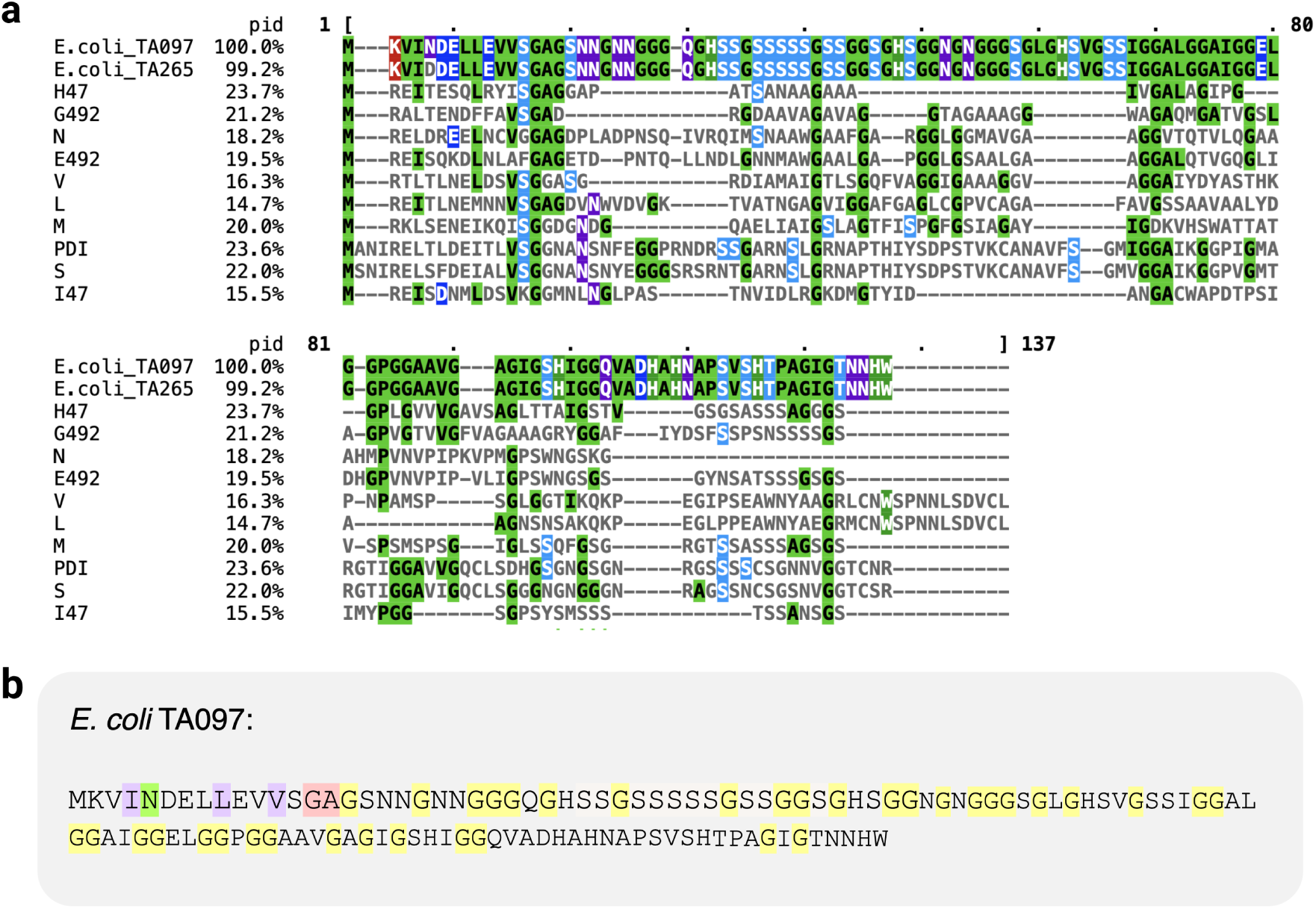
Putative microcins detected within 25 *E. coli* genomes. (a) Alignment of two putative microcins from *E. coli* TA097 and TA265 to the 10 known microcins. “pid” represents percent identity to microcin TA097. (b) Putative microcin sequence TA097. There is a single residue difference between the two found homologs; the other homolog (TA265) contained Aspartic Acid (D) instead of Asparagine (N) at the position highlighted in green. The two residues before the signal sequence cleavage site are shown in pink. Hydrophobic amino acids (purple) are found at positions −4*, −*7, and −12 relative to the cleavage site at the end of the signal sequence. An abundance of glycine (yellow) is found throughout sequence.

**Table 2:** Results from *E. coli*, *Enterobacter*, and *Klebsiella* genomes. In all three sets of genomes, no microcins have previously been detected, while the PCAT microcin exporter gene has been found. The column “unique microcins” refers to the number of unique putative microcins found.

| dataset | genomes | total hits | unique hits | unique microcins |
| --- | --- | --- | --- | --- |
| *E. coli* | 25 | 44 | 20 | 3 |
| *Enterobacter* | 44 | 63 | 26 | 9 |
| *Klebsiella* | 46 | 74 | 19 | 16 |

Next, we used the same method to scan 44 *Enterobacter* spp. and 46 *Klebsiella* spp. genomes from GTDB for potential novel microcins [12]. These two datasets also contained a previously detected PCAT microcin exporter gene but had no detected putative microcins. In total, out of the 44 *Enterobacter* spp. genomes, there were 63 total hits that contained 26 unique sequences (Table 2 and Supplementary File S9). Within these hits, nine had features indicative of class II microcins; they contained a double-glycine signal sequence, had high glycine content, and characteristic hydrophobic residues upstream of the signal sequence (Supplementary File S8). Out of the 46 *Klebsiella* spp. genomes, there were 74 total hits containing a total of 19 unique sequences that we identified as putative microcins (Supplementary File S10). 16 of these hits displayed microcin-like features (Supplementary File S8). However, four of these had a glycine content in the range of 8–10%, which is low compared to known microcins (Supplementary File S6). The other 12 *Klebsiella* spp. sequences had a glycine content of 21–26% and a characteristic double-glycine signal sequence. It is important to note that, while the rate at which microcins were found is good for *Klebsiella* spp. (16/19), only 9/26 *Enterobacter* spp. hits appeared to be true microcins. The reason for this could be that the query embeddings are generated only from the 10 known microcins, eight of which are from *E. coli*, two of which are from *Klebsiella pneumoniae*, and none of which are from *Enterobacter* spp. Therefore, this confirms the need to assess antibacterial activity from newly discovered microcins so that a wider diversity of query embeddings can be used in future studies.

Finally, we looked at 40 additional *E. coli* genomes including 20 from isolates from water and 20 from human extra-intestinal isolates from the B2 phylogroup [25](Supplementary File S5). We chose these particular two genome groups because our prior analysis using cinful identified them as having the least and most class II microcin hits, respectively [12]. For consistency, we limited our comparison to genomes from the B2 phylogroup only. Among the 20 water-sourced genomes, cinful detected a total of eight putative microcins, and our embedding method detected nine (one additional instance of microcin H47 microcin was found by the embeddings and missed by the profile hidden Markov model (pHMM) step in cinful)(Table 3). For the human extra-intestinal genomes, cinful detected 10 putative microcins and the embedding method found 12 (Table 3). The additional two microcins found by the embedding method were also microcins H47 and M, previously missed by the pHMM step of cinful. In summary, out of all 40 genomes analyzed, 18 microcins were found by cinful and 21 were found by the embeddings method. We emphasize, however, that the three additional microcins found by the embedding method were identical at the sequence level to previously known microcins, yet were missed by the pHMM.

**Table 3:** Results for genomes of *E. coli* from the B2 phylogroup isolated from human extra-intestinal and water sources. 15 of the genomes from water-isolated *E. coli* had no previously detected microcins and ten of the genomes from human extra-intestinal-isolated *E. coli* had no previously detected microcins.

| dataset | genomes | cinful hits | embedding hits |
| --- | --- | --- | --- |
| Water | 20 | 8 | 9 |
| Extra-intestinal | 20 | 10 | 12 |

## 4 Discussion

We have assessed the ability of protein large language models, and specifically the embeddings produced by such models, to detect class II microcins in bacterial genomes. We have found that a simple method based on distance in embedding space has much higher sensitivity to discover highly-diverged and relatively short microcin sequences than do sequence-based methods such as BLAST. We have applied our method on a subset of previously analyzed *E. coli*, *Enterobacter* spp., and *Klebsiella* spp. genomes, and have discovered both known and novel putative microcin sequences that prior, sequence-based methods have missed. Our embedding-based method is more computationally costly than a BLAST search, as it requires computation of embeddings for every sequence to be searched against, but it allows the detection of distant, highly diverged, putative homologs that sequence-based methods are likely to miss.

Protein large language models (protein LLMs) have seen massive success in recent years in predicting the biochemical, structural, and sequence properties of proteins [32]. In particular, the popular ESM family of models have been trained on millions of protein sequences and can be used to predict biological structure and function from sequence data [13; 14; 33]. At the core of these models are so-called embeddings, which encode protein characteristics into a high-dimensional numerical space suitable for many different downstream applications. In particular, recent works have shown that embeddings can be used for the identification of protein homologs, even in cases of low sequence similarity [34; 35; 36].

Sequence-based genomic search methods have been the standard for identifying homologous protein sequences. Most of these methods are based on global or local sequence alignments. A critical limitation of sequence-based methods is that they rely on significant sequence similarity between the query and the detected homolog sequence. However, homologs may be diverged in sequence while retaining their overall structure and function [34]. Such is the case with class II microcins; they are a unique group of proteins that are both very small and have high sequence divergence relative to each other.

The short length of class II microcins creates problems for sequence-based methods but provides advantages for the embedding method. First, it reduces the computational cost of generating and storing the embeddings for a sequence; since each residue in a protein sequence produces an embedding, the shorter the sequence, the fewer embeddings need to be generated. Secondly, a shorter sequence reduces the “washing out” effect of encoded features due to averaging of embeddings. In other words, the longer the protein, the more likely that specific features will be lost due to a regression-to-the-mean effect. Finally, class II microcins also have high sequence divergence, which provides additional challenges for alignment-based methods. This makes class II microcins great candidates for a semantic homology search while outperforming the canonical alignment-based search methods.

The known microcins have several characteristics that distinguish them from non-microcin peptides, including specific features of their signal sequences, glycine content, and genomic location near their PCAT exporter [7]. Our embedding method recovered novel sequences that unequivocally displayed these characteristics (Table 2). Additionally, however, we also found hits that do not fully look like canonical microcins (Supplementary File S6). In particular, some of our detected hits had only around 8–10% glycine content, yet displayed other microcin-like features such as a double-glycine signal, appropriate hydrophobic residues upstream of the signal, and a microcin-like size (*<* 150 residues). These sequences are interesting candidates for future experimental validation.

We would like to emphasize that some of the hits we identified did not appear to have microcin-like features and are likely false positives. Many of these false positive did have a high glycine content, which may explain why they were found by the embeddings. However, they lacked other microcin features, such as a double-glycine signal sequence. As mentioned above, one reason why false positives may be identified is that we are using averaged embeddings, and certain important protein features may be lost during the averaging step. A second reason for false positives is that our method for identifying microcin hits relies on a simple distance metric and distance cut-off; therefore, we can expect to occasionally find sequences that are not microcins, but may share some characteristics with real microcins. These peptides could have either lost some of their microcin properties over evolutionary time or by coincidence just have microcin-like features such as high glycine content. Although we found apparent non-microcin hits, our method has very good sensitivity and does not appear to miss microcins when they are present in the genome.

Our work has also revealed that our prior tool for microcin search, cinful [12], can miss both known microcins and novel putative microcin sequences. We found candidate sequences to be lost at all three stages of cinful: the ORF extraction stage via Prodigal [31], the BLAST stage, and the profile hidden Markov model (pHMM) stage. Because here we used an extremely broad definition of ORF, considering all sequences in the microcin length range starting with a start codon and ending with a stop codon, our ORF extraction step was unlikely to overlook any potential microcins. It is worth noting that we did not use alternative start codons because it would have vastly inflated the number of extracted ORFs for each assembly. However, since alternative start codons are rare, we believe this causes minimal problems in detecting microcins. Next, as shown here, BLAST tends to have difficulty identifying microcins because they are short and highly diverged. In addition, BLAST tends to generate false positives, i.e., it recovers sequences which are not microcins. This propensity of BLAST to generate false positives required the addition of a final filtering stage, the pHMM stage, in cinful. But generating a good pHMM was hindered by the small number of known microcins (10) [12]. As more microcins get discovered and verified, it is likely that both BLAST and pHMM searches can become more accurate, and an updated future version of cinful with a larger set of reference microcin sequences may perform better.

Our ORF extraction approach was extremely exhaustive, thus minimizing the chance of missed hits. However, this approach comes with the drawback of substantially increased computational cost, as all the putative ORFs need to be processed, and embedding calculations are not cheap. We believe that in the future, a more targeted search strategy may be appropriate and sufficient. Microcins tend to co-locate with their (highly conserved) peptidase-containing ABC transporter (PCAT) and membrane fusion protein (MFP), which are used for their export. Because PCATs and MFPs are so conserved, they are easier to identify in a sequence-based search than are microcins [7; 12]. Therefore, future searches could restrict the search space only to ORFs located within close proximity to PCAT or MFP sequences.

In summary, our work has shown that protein embeddings produced by large language models can be used successfully to search for small, highly diverged sequences such as class II microcins in genome-scale datasets. Embeddings have several advantaged over sequence-based methods and appear to have both higher sensitivity and higher specificity for detecting novel microcins. However, embeddings incur substantially higher computational costs than a BLAST search or a hidden Markov model and therefore may only be suitable in cases where sequence-based methods clearly fail or where the search space is somewhat limited.

## Acknowledgments

This work was funded by the National Institute of Allergy and Infectious Diseases of the National Institutes of Health (award numbers R01 AI148419 and R21 AI159203 to B.W.D.), Tito’s Handmade Vodka (B.W.D.), the Blumberg Centennial Professorship in Molecular Evolution (C.O.W.), and the Reeder Centennial Fellowship in Systematic and Evolutionary Biology at The University of Texas at Austin (C.O.W.).

