## Supplementary figures and supplementary file captions for "Semantic search using protein large language models detects class II microcins in bacterial genomes"

### Supplementary Figures and Files

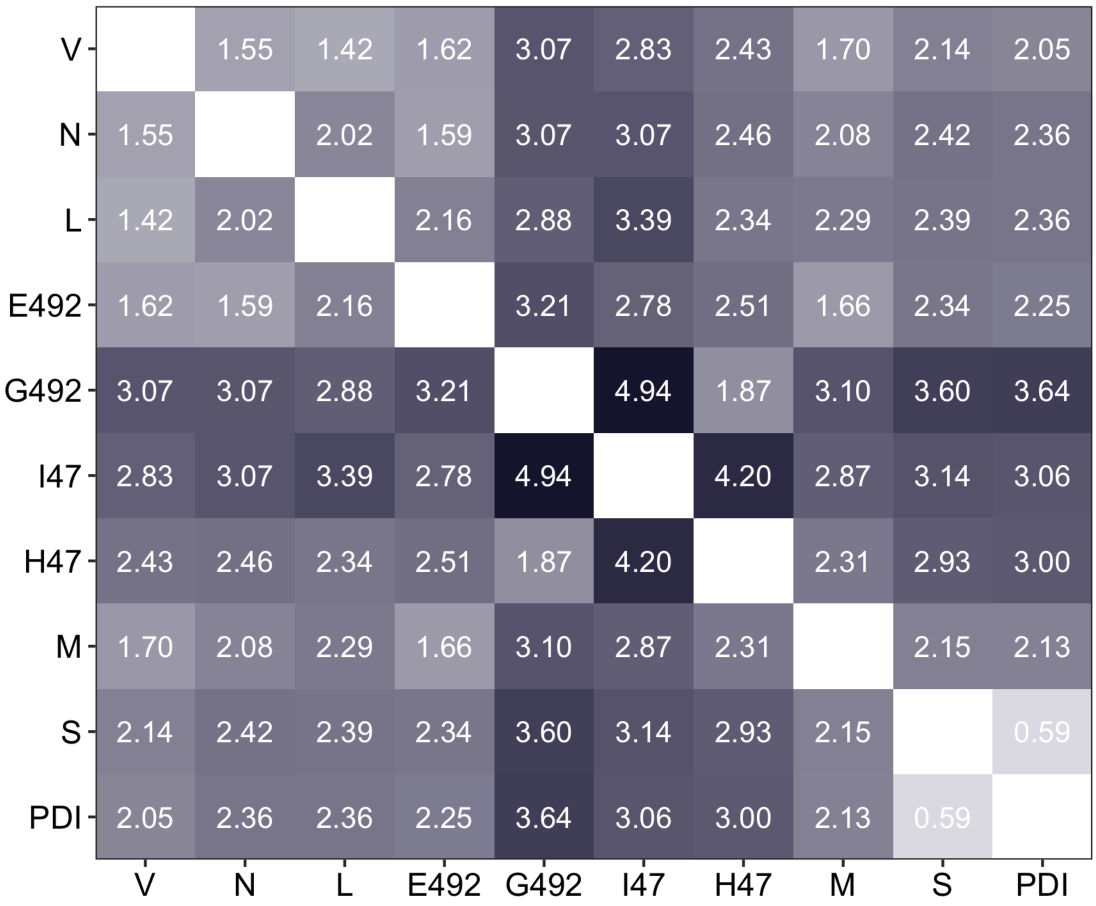


**Supplementary Figure S1:** Embedding distance between all pairs of the ten known microcins.

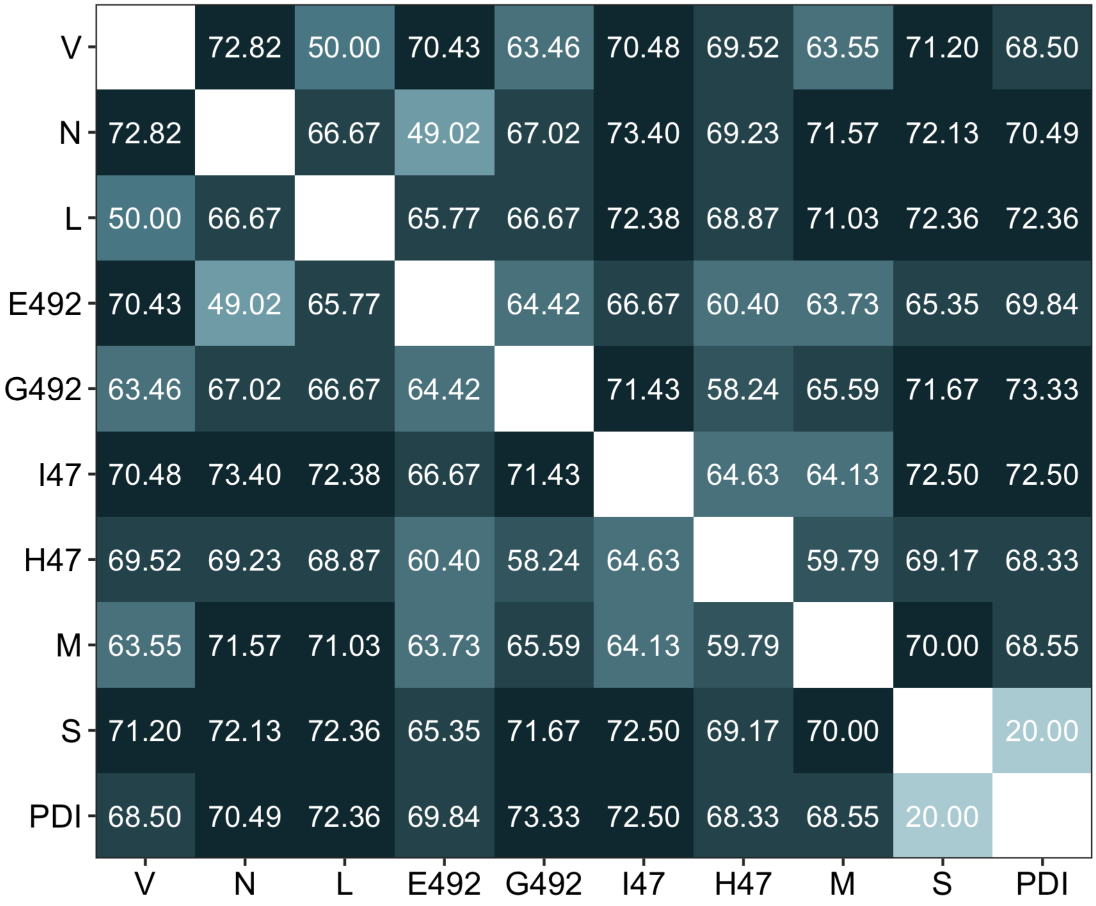


**Supplementary Figure S2:** Percent sequence divergence between all pairs of the ten known microcins.

**Supplementary File S1:** Information about the dataset of 25 *E. coli* genomes. Contains the columns `name`, `strain_name`, `accenssion_number`, `strain_category`, `phylogroup`, `Genome_ID`, `microcin_hit_count`, `CvaB_hit_count`.

**Supplementary File S2:** Information about the dataset of *Enterobacter* genomes. Contains the columns `accession`, `ncbi_organism_name`, `cinful_CvaB_found` (indicates whether a CvaB was previously found by cinful—the value is `TRUE` for all genomes in this set), `cinful_microcins_found`.

**Supplementary File S3:** Information about the dataset of *Klebsiella* genomes. Contains the columns `accession`, `ncbi_organism_name`, `cinful_microcins_found` (number of microcins found by cinful), `cinful_CvaB_found` (indicates whether a CvaB was previously found by cinful—the value is `TRUE` for all genomes in this set).

**Supplementary File S4:** Cinful microcin and CvaB hits from the Touchon dataset. Contains the columns `name`, `strain_name`, `accession_number`, `strain_category` (origin of the bacteria), `phylogroup`, `Genome_ID`, `microcin_hit_count` (how many microcins were detected by Cinful), `CvaB_hit_count` (the number of microcin exporter proteins Cinful detected), `Assembly` (accession number for retrieval from NCBI).

**Supplementary File S5:** Cinful results for 40 water-sourced and extra-intestinal genomes from the Touchon dataset. Contains the columns `name`, `strain_name`, `accession_number`, `strain_category` (origin of the bacteria), `phylogroup`, `Genome_ID`, `microcin_hit_count` (how many microcins were detected by Cinful), `CvaB_hit_count` (if Cinful detected a microcin exporter protein), `Assembly` (accession number for retrieval from NCBI).

**Supplementary File S6:** Amino acid sequences of the ten known microcins, provided in FASTA format.

**Supplementary File S7:** Alignment of embedding hits collected from 25 *E. coli* genomes.

**Supplementary File S8:** Putative microcins found in the *E. coli*, *Enterobacter*, and *Klebsiella* datasets.

**Supplementary File S9:** Alignment of embedding hits collected from 44 *Enterobacter* genomes.

**Supplementary File S10:** Alignment of embedding hits collected from 46 *Klebsiella* genomes.
