## Supplementary File S7 for "Semantic search using protein large language models detects class II microcins in bacterial genomes"

```
Reference sequence (1): GCA_000240325.1_ORF.81043
Identities normalised by aligned length.
Colored by: identity

                                         cov    pid   1 [         .         .         .         .         :         .         .         . 80
 1 GCA_000240325.1_ORF.81043         100.0% 100.0%     MKQLTVFEMEEISGGYSWDFSSVANALSSIICNGAEAIGASILGATAGALAGSAMGGIFG-GNGGGILGIGSIGQAVGMI
 2 JAND01000096.1_ORF.89371          100.0% 100.0%     MKQLTVFEMEEISGGYSWDFSSVANALSSIICNGAEAIGASILGATAGALAGSAMGGIFG-GNGGGILGIGSIGQAVGMI
 3 CP004887.1_ORF.29173              100.0% 100.0%     MKQLTVFEMEEISGGYSWDFSSVANALSSIICNGAEAIGASILGATAGALAGSAMGGIFG-GNGGGILGIGSIGQAVGMI
 4 JUYF01000041.1_ORF.80449          100.0% 100.0%     MKQLTVFEMEEISGGYSWDFSSVANALSSIICNGAEAIGASILGATAGALAGSAMGGIFG-GNGGGILGIGSIGQAVGMI
 5 CP017450.1_ORF.58218              100.0% 100.0%     MKQLTVFEMEEISGGYSWDFSSVANALSSIICNGAEAIGASILGATAGALAGSAMGGIFG-GNGGGILGIGSIGQAVGMI
 6 JAKW01000011.1_ORF.7894           100.0% 100.0%     MKQLTVFEMEEISGGYSWDFSSVANALSSIICNGAEAIGASILGATAGALAGSAMGGIFG-GNGGGILGIGSIGQAVGMI
 7 VOJB01000088.1_ORF.64837          100.0% 100.0%     MKQLTVFEMEEISGGYSWDFSSVANALSSIICNGAEAIGASILGATAGALAGSAMGGIFG-GNGGGILGIGSIGQAVGMI
 8 WCIN01000037.1_ORF.41825          100.0% 100.0%     MKQLTVFEMEEISGGYSWDFSSVANALSSIICNGAEAIGASILGATAGALAGSAMGGIFG-GNGGGILGIGSIGQAVGMI
 9 VLLY01000008.1_ORF.86873          100.0% 100.0%     MKQLTVFEMEEISGGYSWDFSSVANALSSIICNGAEAIGASILGATAGALAGSAMGGIFG-GNGGGILGIGSIGQAVGMI
10 FKYS01000010.1_ORF.43257          100.0% 100.0%     MKQLTVFEMEEISGGYSWDFSSVANALSSIICNGAEAIGASILGATAGALAGSAMGGIFG-GNGGGILGIGSIGQAVGMI
11 ALNJ01000099.1_ORF.64562          100.0%  94.2%     MKQLTVFEMEEISGGYSWDFSSITGALTSIVFNSAEAIGASILGATAGALAGSAMGGIFG-GNGGGILGIGSIGQAVGMI
12 JAGZTV010000025.1_ORF.59009       100.0%  94.2%     MKQLTVFEMEEISGGYSWDFSSITGALTSIVFNSAEAIGASILGATAGALAGSAMGGIFG-GNGGGILGIGSIGQAVGMI
13 CABGKN010000008.1_ORF.45398       100.0%  94.2%     MKQLTVFEMEEISGGYSWDFSSITGALTSIVFNSAEAIGASILGATAGALAGSAMGGIFG-GNGGGILGIGSIGQAVGMI
14 CP020358.1_ORF.82951              100.0%  93.3%     MKQLTVFEMEEISGGYCWDFSSITGALTSIVFNSAEAIGASILGATAGALAGSAMGGIFG-GNGGGILGIGSIGQAVGMI
15 CABGII010000013.1_ORF.42080       100.0%  93.3%     MKQLTVFEMEEISGGYCWDFSSITGALTSIVFNSAEAIGASILGATAGALAGSAMGGIFG-GNGGGILGIGSIGQAVGMI
16 CP008788.1_ORF.73006              100.0%  95.0%     MKQLTVFEMEEISGGYSWDFSSVANALSSIICNGAEAIGASILGATAGALAGSAMGGIFG-GNGGGILGIGSIGQAVGMI
17 PQKN01000027.1_ORF.44266          100.0%  95.0%     MKQLTVFEMEEISGGYSWDFSSVANALSSIICNGAEAIGASILGATAGALAGSAMGGIFG-GNGGGILGIGSIGQAVGMI
18 PQKM01000026.1_ORF.46098          100.0%  95.0%     MKQLTVFEMEEISGGYSWDFSSVANALSSIICNGAEAIGASILGATAGALAGSAMGGIFG-GNGGGILGIGSIGQAVGMI
19 JAERPV010000004.1_ORF.67367       100.0%  95.0%     MKQLTVFEMEEISGGYSWDFSSVANALSSIICNGAEAIGASILGATAGALAGSAMGGIFG-GNGGGILGIGSIGQAVGMI
20 KI535597.1_ORF.26878              100.0%  58.7%     MKQLTVFEMEEISGGYSWDFSSLGNALSSMAGNAVELVASVAVAASAGGMAGSVIGGRWG-GAGGGILGFGAIGQGVGMI
21 KI535631.1_ORF.84463              100.0%  58.7%     MKQLTVFEMEEISGGYSWDFSSLGNALSSMAGNAVELVASVAVAASAGGMAGSVIGGRWG-GAGGGILGFGAIGQGVGMI
22 JAKY01000009.1_ORF.7757           100.0%  58.7%     MKQLTVFEMEEISGGYSWDFSSLGNALSSMAGNAVELVASVAVAASAGGMAGSVIGGRWG-GAGGGILGFGAIGQGVGMI
23 JAAFEW010000005.1_ORF.74831       100.0%  58.7%     MKQLTVFEMEEISGGYSWDFSSLGNALSSMAGNAVELVASVAVAASAGGMAGSVIGGRWG-GAGGGILGFGAIGQGVGMI
24 FKYZ01000010.1_ORF.52325          100.0%  58.7%     MKQLTVFEMEEISGGYSWDFSSLGNALSSMAGNAVELVASVAVAASAGGMAGSVIGGRWG-GAGGGILGFGAIGQGVGMI
25 FKZZ01000010.1_ORF.54370          100.0%  58.7%     MKQLTVFEMEEISGGYSWDFSSLGNALSSMAGNAVELVASVAVAASAGGMAGSVIGGRWG-GAGGGILGFGAIGQGVGMI
26 ARVT01000001.1_ORF.71550          100.0%  53.7%     MKQLTVFEMEAISGGYSWDMSSIGGILTSLVSNAAELATSATLGASVGGIYGSIIGGRWG-GAGGGILGFGAIGQGVGMV
27 JAKX01000046.1_ORF.49662          100.0%  53.7%     MKQLTVFEMEAISGGYSWDMSSIGGILTSLVSNAAELATSATLGASVGGIYGSIIGGRWG-GAGGGILGFGAIGQGVGMV
28 KK097710.1_ORF.57938              100.0%  53.7%     MKQLTVFEMEAISGGYSWDMSSIGGILTSLVSNAAELATSATLGASVGGIYGSIIGGRWG-GAGGGILGFGAIGQGVGMV
29 KQ235791.1_ORF.65204              100.0%  53.7%     MKQLTVFEMEAISGGYSWDMSSIGGILTSLVSNAAELATSATLGASVGGIYGSIIGGRWG-GAGGGILGFGAIGQGVGMV
30 BCZK01000008.1_ORF.48858          100.0%  53.7%     MKQLTVFEMEAISGGYSWDMSSIGGILTSLVSNAAELATSATLGASVGGIYGSIIGGRWG-GAGGGILGFGAIGQGVGMV
31 DIEF01000027.1_ORF.35923          100.0%  53.7%     MKQLTVFEMEAISGGYSWDMSSIGGILTSLVSNAAELATSATLGASVGGIYGSIIGGRWG-GAGGGILGFGAIGQGVGMV
32 LR890312.1_ORF.42131              100.0%  53.7%     MKQLTVFEMEAISGGYSWDMSSIGGILTSLVSNAAELATSATLGASVGGIYGSIIGGRWG-GAGGGILGFGAIGQGVGMV
33 AKCF01000001.1_ORF.31003          100.0%  52.5%     MKQLTVFEMEEISGGYSWDFSSIQSTISSFACNAVEAIGAGVLGGIVGGSFGTLIGGTQS-GANGGLLGFGLIGNGVGLV
34 GCA_014169355_ORF.72421            97.5%  33.1%     MKQLTIVEMNDVSGA---GIQQIIDGSQNVLFGLLDAAIGAVLLGFGASGAGGLQGGVTGNGSGGGILGFGVITIGIGSI
35 CP039791.1_ORF.23972               97.5%  33.1%     MKQLTIVEMNDVSGA---GIQQIIDGSQNVLFGLLDAAIGAVLLGFGASGAGGLQGGVTGNGSGGGILGFGVITIGIGSI
36 GCA_014189245.1_ORF.23815          97.5%  31.5%     MKELTTIEMNDISGA---GIKQILDGSQNMFAGLVDATVGAAIGLIGYASGGGLQGGLTGNGSGGGLLGFGTIAMGIGAL
37 GCA_014189245.1_ORF.23816          97.5%  30.5%     MKELTAIEMDDISGA---GIRQILDGTQDLLSGLVDATVGATIGLIGYASGGGLQGGLTGNGAGGGALGFGTIAMGIGAL
38 WMOU01000014.1_ORF.12806           97.5%  33.9%     MKELTSIEMNDVSGA---GIKDMINGSQSFLSGALDTVLGAALGAIAYSSIFGLQGGVTGNGSGGGIIGLGIITTAVGSI
39 JAFHNU010000005.1_ORF.70092        97.5%  33.9%     MKELTSIEMNDVSGA---GIKDMINGSQSFLSGALDTVLGAALGAIAYSSIFGLQGGVTGNGSGGGIIGLGIITTAVGSI
40 JAFHNV010000002.1_ORF.38562        97.5%  33.9%     MKELTSIEMNDVSGA---GIKDMINGSQSFLSGALDTVLGAALGAIAYSSIFGLQGGVTGNGSGGGIIGLGIITTAVGSI
41 JAFHNS010000005.1_ORF.69110        97.5%  33.9%     MKELTSIEMNDVSGA---GIKDMINGSQSFLSGALDTVLGAALGAIAYSSIFGLQGGVTGNGSGGGIIGLGIITTAVGSI
42 JAFHWD010000004.1_ORF.66056        97.5%  33.9%     MKELTSIEMNDVSGA---GIKDMINGSQSFLSGALDTVLGAALGAIAYSSIFGLQGGVTGNGSGGGIIGLGIITTAVGSI
43 JAFHWF010000004.1_ORF.64614        97.5%  33.9%     MKELTSIEMNDVSGA---GIKDMINGSQSFLSGALDTVLGAALGAIAYSSIFGLQGGVTGNGSGGGIIGLGIITTAVGSI
44 JAFHUV010000004.1_ORF.64984        97.5%  33.9%     MKELTSIEMNDVSGA---GIKDMINGSQSFLSGALDTVLGAALGAIAYSSIFGLQGGVTGNGSGGGIIGLGIITTAVGSI
45 CABGYN010000032.1_ORF.68677        97.5%  33.9%     MKELTSIEMNDVSGA---GIKDMINGSQSFLSGALDTVLGAALGAIAYSSIFGLQGGVTGNGSGGGIIGLGIITTAVGSI
46 GCA_000240325.1_ORF.64824          79.2%  18.0%     MKELTIGEMESISGGF--NLLGFANSITSFIVDNGNYLSDFITSA----------GATIANAIVNDTVEFAKFLTGASDW
47 JAND01000047.1_ORF.50276           79.2%  18.0%     MKELTIGEMESISGGF--NLLGFANSITSFIVDNGNYLSDFITSA----------GATIANAIVNDTVEFAKFLTGASDW
48 CP004887.1_ORF.41180               79.2%  18.0%     MKELTIGEMESISGGF--NLLGFANSITSFIVDNGNYLSDFITSA----------GATIANAIVNDTVEFAKFLTGASDW
49 CP008788.1_ORF.70925               79.2%  18.0%     MKELTIGEMESISGGF--NLLGFANSITSFIVDNGNYLSDFITSA----------GATIANAIVNDTVEFAKFLTGASDW
50 JUYF01000502.1_ORF.13288           79.2%  18.0%     MKELTIGEMESISGGF--NLLGFANSITSFIVDNGNYLSDFITSA----------GATIANAIVNDTVEFAKFLTGASDW
51 CP017450.1_ORF.56285               79.2%  18.0%     MKELTIGEMESISGGF--NLLGFANSITSFIVDNGNYLSDFITSA----------GATIANAIVNDTVEFAKFLTGASDW
52 JAKW01000015.1_ORF.54652           79.2%  18.0%     MKELTIGEMESISGGF--NLLGFANSITSFIVDNGNYLSDFITSA----------GATIANAIVNDTVEFAKFLTGASDW
53 PQKN01000002.1_ORF.29788           79.2%  18.0%     MKELTIGEMESISGGF--NLLGFANSITSFIVDNGNYLSDFITSA----------GATIANAIVNDTVEFAKFLTGASDW
54 PQKM01000001.1_ORF.2990            79.2%  18.0%     MKELTIGEMESISGGF--NLLGFANSITSFIVDNGNYLSDFITSA----------GATIANAIVNDTVEFAKFLTGASDW
55 VOJB01000079.1_ORF.54930           79.2%  18.0%     MKELTIGEMESISGGF--NLLGFANSITSFIVDNGNYLSDFITSA----------GATIANAIVNDTVEFAKFLTGASDW
56 WCIN01000041.1_ORF.61881           79.2%  18.0%     MKELTIGEMESISGGF--NLLGFANSITSFIVDNGNYLSDFITSA----------GATIANAIVNDTVEFAKFLTGASDW
57 VLLY01000011.1_ORF.8219            79.2%  18.0%     MKELTIGEMESISGGF--NLLGFANSITSFIVDNGNYLSDFITSA----------GATIANAIVNDTVEFAKFLTGASDW
58 JADRTN010000001.1_ORF.648          79.2%  18.0%     MKELTIGEMESISGGF--NLLGFANSITSFIVDNGNYLSDFITSA----------GATIANAIVNDTVEFAKFLTGASDW
59 ALNJ01000086.1_ORF.55451           79.2%  20.7%     MKELTIGEMESISGGF--NLFGFANSITSLIINSGNHLSDFITSA----------GATIANAIVNGTVEFGKFLTGASDW
60 CP020358.1_ORF.49881               79.2%  20.7%     MKELTIGEMESISGGF--NLFGFANSITSLITNSGNHLSDFITSA----------GATIANAVVNGTVEFGKFLTGASDW
61 AKCF01000001.1_ORF.13999           79.2%  18.3%     MKELTQYEMENISGA----------GLFDLPCALVDFTIQSALGLMKVGFDGSLTIAKTALETSLDML--SNLLSGSPSS
62 KI535631.1_ORF.67631               79.2%  18.3%     MKELTQYEMENISGA----------GLFDLPCALVDFTIQSALGLMKVGFDGSLTIAKTALETSLDML--SNLLSGSPSS
63 ARVT01000001.1_ORF.83832           79.2%  18.3%     MKELTQYEMENISGA----------GLFDLPCALVDFTIQSALGLMKVGFDGSLTIAKTALETSLDML--SNLLSGSPSS
64 JAKX01000001.1_ORF.726             79.2%  18.3%     MKELTQYEMENISGA----------GLFDLPCALVDFTIQSALGLMKVGFDGSLTIAKTALETSLDML--SNLLSGSPSS
65 KK097709.1_ORF.46887               79.2%  18.3%     MKELTQYEMENISGA----------GLFDLPCALVDFTIQSALGLMKVGFDGSLTIAKTALETSLDML--SNLLSGSPSS
66 BCZK01000002.1_ORF.16754           79.2%  18.3%     MKELTQYEMENISGA----------GLFDLPCALVDFTIQSALGLMKVGFDGSLTIAKTALETSLDML--SNLLSGSPSS
67 DIEF01000003.1_ORF.23844           79.2%  18.3%     MKELTQYEMENISGA----------GLFDLPCALVDFTIQSALGLMKVGFDGSLTIAKTALETSLDML--SNLLSGSPSS
68 KI535597.1_ORF.31267               79.2%  18.3%     MKELTQYEMENISGA----------GLFDLPCALVDFTIQSALGLMKVGFDGSLTIAKTALETSLDML--SNLLSGSPSS
69 JAKY01000041.1_ORF.44918           79.2%  18.3%     MKELTQYEMENISGA----------GLFDLPCALVDFTIQSALGLMKVGFDGSLTIAKTALETSLDML--SNLLSGSPSS
70 KQ235791.1_ORF.67988               79.2%  18.3%     MKELTQYEMENISGA----------GLFDLPCALVDFTIQSALGLMKVGFDGSLTIAKTALETSLDML--SNLLSGSPSS
71 JAND01000082.1_ORF.77829           76.7%  21.0%     MT--------TICPQC-----------------GSDRVSERNIGRKAGGIAGGVAGALSGAGTGAAI---GSVVPVVGTA
72 JADRTN010000013.1_ORF.51264        76.7%  21.0%     MT--------TICPQC-----------------GSDRVSERNIGRKAGGIAGGVAGALSGAGTGAAI---GSVVPVVGTA
73 AP022142.1_ORF.50706               76.7%  21.0%     MT--------TICPQC-----------------GSDRVSERNIGRKAGGIAGGVAGALSGAGTGAAI---GSVVPVVGTA
74 AP022142.1_ORF.3785                76.7%  21.8%     MT--------TICPEC-----------------GSDRVSERNIGRKAGSIAGGVAGALSGAGTGAAI---GSVVPVLGTA
   consensus/100%                                      Mp........tlsst....................phh.t..h.h..........huh.u.t.s.sh...u.hh.s.s..
   consensus/90%                                       MKpLT.hEMpplSGu.............sh..s.hphh.t.hhuh..........huhhu.t.s.shl..u.hh.uss..
   consensus/80%                                       MKpLT.hEMppISGu...sh.th.su.pshhssssphh.s.hhut..........Guhhu.usssshlthuthhpusu.h
   consensus/70%                                       MKpLThhEMEpISGuh..sh.shhsulpShlssss-hlssshluA.hhu..huh.Guhhu.ususshlthutlhpusu.h

                                         cov    pid  81         .         1         .         .         .         . ] 143
 1 GCA_000240325.1_ORF.81043         100.0% 100.0%     YGLCVG-----------GVGG-AIAGAMVGWDITLDKCIEMFDGFVDGTLAYW----------
 2 JAND01000096.1_ORF.89371          100.0% 100.0%     YGLCVG-----------GVGG-AIAGAMVGWDITLDKCIEMFDGFVDGTLAYW----------
 3 CP004887.1_ORF.29173              100.0% 100.0%     YGLCVG-----------GVGG-AIAGAMVGWDITLDKCIEMFDGFVDGTLAYW----------
 4 JUYF01000041.1_ORF.80449          100.0% 100.0%     YGLCVG-----------GVGG-AIAGAMVGWDITLDKCIEMFDGFVDGTLAYW----------
 5 CP017450.1_ORF.58218              100.0% 100.0%     YGLCVG-----------GVGG-AIAGAMVGWDITLDKCIEMFDGFVDGTLAYW----------
```

```
 6 JAKW01000011.1_ORF.7894      100.0% 100.0%   YGLCVG-----------GVGG-AIAGAMVGWDITLDKCIEMFDGFVDGTLAYW----------
 7 VOJB01000088.1_ORF.64837     100.0% 100.0%   YGLCVG-----------GVGG-AIAGAMVGWDITLDKCIEMFDGFVDGTLAYW----------
 8 WCIN01000037.1_ORF.41825     100.0% 100.0%   YGLCVG-----------GVGG-AIAGAMVGWDITLDKCIEMFDGFVDGTLAYW----------
 9 VLLY01000008.1_ORF.86873     100.0% 100.0%   YGLCVG-----------GVGG-AIAGAMVGWDITLDKCIEMFDGFVDGTLAYW----------
10 FKYS01000010.1_ORF.43257     100.0% 100.0%   YGLCVG-----------GVGG-AIAGAMVGWDITLDKCIEMFDGFVDGTLAYW----------
11 ALNJ01000099.1_ORF.64562     100.0%  94.2%   YGLCVG-----------GVGG-AIAGAMVGWDITLDKCIEMFDGFVDGTLAYW----------
12 JAGZTV010000025.1_ORF.59009  100.0%  94.2%   YGLCVG-----------GVGG-AIAGAMVGWDITLDKCIEMFDGFVDGTLAYW----------
13 CABGKN010000008.1_ORF.45398  100.0%  94.2%   YGLCVG-----------GVGG-AIAGAMVGWDITLDKCIEMFDGFVDGTLAYW----------
14 CP020358.1_ORF.82951         100.0%  93.3%   YGLCVG-----------GVGG-AIAGAMVGWDITLDKCIEMFDGFVDGTLAYW----------
15 CABGII010000013.1_ORF.42080  100.0%  93.3%   YGLCVG-----------GVGG-AIAGAMVGWDITLDKCIEMFDGFVDGTLAYW----------
16 CP008788.1_ORF.73006         100.0%  95.0%   CGLYIG-----------GLGG-AVAGVMVGWDITLDKCIEMFDGFVDGTLAYW----------
17 PQKN01000027.1_ORF.44266     100.0%  95.0%   CGLYIG-----------GLGG-AVAGVMVGWDITLDKCIEMFDGFVDGTLAYW----------
18 PQKM01000026.1_ORF.46098     100.0%  95.0%   CGLYIG-----------GLGG-AVAGVMVGWDITLDKCIEMFDGFVDGTLAYW----------
19 JAERPV010000004.1_ORF.67367  100.0%  95.0%   CGLYIG-----------GLGG-AVAGVMVGWDITLDKCIEMFDGFVDGTLAYW----------
20 KI535597.1_ORF.26878         100.0%  58.7%   WGLVVG-----------AIGC-GIAGAFVGWDVVSTEAIALAEGVINGTTKLWS---------
21 KI535631.1_ORF.84463         100.0%  58.7%   WGLVVG-----------AIGC-GIAGAFVGWDVVSTEAIALAEGVINGTTKLWS---------
22 JAKY01000009.1_ORF.7757      100.0%  58.7%   WGLVVG-----------AIGC-GIAGAFVGWDVVSTEAIALAEGVINGTTKLWS---------
23 JAAFEW010000005.1_ORF.74831  100.0%  58.7%   WGLVVG-----------AIGC-GIAGAFVGWDVVSTEAIALAEGVINGTTKLWS---------
24 FKYZ01000010.1_ORF.52325     100.0%  58.7%   WGLVVG-----------AIGC-GIAGAFVGWDVVSTEAIALAEGVINGTTKLWS---------
25 FKZZ01000010.1_ORF.54370     100.0%  58.7%   WGLVVG-----------AIGC-GIAGAFVGWDVVSTEAIALAEGVINGTTKLWS---------
26 ARVT01000001.1_ORF.71550     100.0%  53.7%   WGLVVG-----------AIGC-GVAASVVGWDKTYELAMGAIAGSIDGTLTPWN---------
27 JAKX01000046.1_ORF.49662     100.0%  53.7%   WGLVVG-----------AIGC-GVAASVVGWDKTYELAMGAIAGSIDGTLTPWN---------
28 KK097710.1_ORF.57938         100.0%  53.7%   WGLVVG-----------AIGC-GVAASVVGWDKTYELAMGAIAGSIDGTLTPWN---------
29 KQ235791.1_ORF.65204         100.0%  53.7%   WGLVVG-----------AIGC-GVAASVVGWDKTYELAMGAIAGSIDGTLTPWN---------
30 BCZK01000008.1_ORF.48858     100.0%  53.7%   WGLVVG-----------AIGC-GVAASVVGWDKTYELAMGAIAGSIDGTLTPWN---------
31 DIEF01000027.1_ORF.35923     100.0%  53.7%   WGLVVG-----------AIGC-GVAASVVGWDKTYELAMGAIAGSIDGTLTPWN---------
32 LR890312.1_ORF.42131         100.0%  53.7%   WGLVVG-----------AIGC-GVAASVVGWDKTYELAMGAIAGSIDGTLTPWN---------
33 AKCF01000001.1_ORF.31003     100.0%  52.5%   WGAIVG-----------GVAG-ASGAIAAGWDTTLQVVVDAFQSVIDGTFIPWSH--------
34 GCA_014169355_ORF.72421       97.5%  33.1%   WGAIQG-----------AVTG-AVWGAYVGADTSVEYIKKGVDAWFAGTIGGWTPN-------
35 CP039791.1_ORF.23972          97.5%  33.1%   WGAIQG-----------AVTG-AVWGAYVGADTSVEYIKKGVDAWFAGTIGGWTPN-------
36 GCA_014189245.1_ORF.23815     97.5%  31.5%   WGAIQG-----------GIFG-TIMGAYNGADYVNGQITRLIDGILDGTAGGFKAN-------
37 GCA_014189245.1_ORF.23816     97.5%  30.5%   WGAIQG-----------GIFG-AIMGAYNGADYVNGQVTRLIDGILDGTAGGFKANGGFSISF
38 WMOU01000014.1_ORF.12806      97.5%  33.9%   WGAIQG-----------AVWG-GMQGAYNGADYINGQVTDMINGIIDGTAGGFSSK-------
39 JAFHNU010000005.1_ORF.70092   97.5%  33.9%   WGAIQG-----------AVWG-GMQGAYNGADYINGQVTDMINGIIDGTAGGFSSK-------
40 JAFHNV010000002.1_ORF.38562   97.5%  33.9%   WGAIQG-----------AVWG-GMQGAYNGADYINGQVTDMINGIIDGTAGGFSSK-------
41 JAFHNS010000005.1_ORF.69110   97.5%  33.9%   WGAIQG-----------AVWG-GMQGAYNGADYINGQVTDMINGIIDGTAGGFSSK-------
42 JAFHWD010000004.1_ORF.66056   97.5%  33.9%   WGAIQG-----------AVWG-GMQGAYNGADYINGQVTDMINGIIDGTAGGFSSK-------
43 JAFHWF010000004.1_ORF.64614   97.5%  33.9%   WGAIQG-----------AVWG-GMQGAYNGADYINGQVTDMINGIIDGTAGGFSSK-------
44 JAFHUV010000004.1_ORF.64984   97.5%  33.9%   WGAIQG-----------AVWG-GMQGAYNGADYINGQVTDMINGIIDGTAGGFSSK-------
45 CABGYN010000032.1_ORF.68677   97.5%  33.9%   WGAIQG-----------AVWG-GMQGAYNGADYINGQVTDMINGIIDGTAGGFSSK-------
46 GCA_000240325.1_ORF.64824     79.2%  18.0%   ENYVAA---------SNENWSNAVHNLSGEWNTFTNSITA-----------------------
47 JAND01000047.1_ORF.50276      79.2%  18.0%   ENYVAA---------SNENWSNAVHNLSGEWNTFTNSITA-----------------------
48 CP004887.1_ORF.41180          79.2%  18.0%   ENYVAA---------SNENWSNAVHNLSGEWNTFTNSITA-----------------------
49 CP008788.1_ORF.70925          79.2%  18.0%   ENYVAA---------SNENWSNAVHNLSGEWNTFTNSITA-----------------------
50 JUYF01000502.1_ORF.13288      79.2%  18.0%   ENYVAA---------SNENWSNAVHNLSGEWNTFTNSITA-----------------------
51 CP017450.1_ORF.56285          79.2%  18.0%   ENYVAA---------SNENWSNAVHNLSGEWNTFTNSITA-----------------------
52 JAKW01000015.1_ORF.54652      79.2%  18.0%   ENYVAA---------SNENWSNAVHNLSGEWNTFTNSITA-----------------------
53 PQKN01000002.1_ORF.29788      79.2%  18.0%   ENYVAA---------SNENWSNAVHNLSGEWNTFTNSITA-----------------------
54 PQKM01000001.1_ORF.2990       79.2%  18.0%   ENYVAA---------SNENWSNAVHNLSGEWNTFTNSITA-----------------------
55 VOJB01000079.1_ORF.54930      79.2%  18.0%   ENYVAA---------SNENWSNAVHNLSGEWNTFTNSITA-----------------------
56 WCIN01000041.1_ORF.61881      79.2%  18.0%   ENYVAA---------SNENWSNAVHNLSGEWNTFTNSITA-----------------------
57 VLLY01000011.1_ORF.8219       79.2%  18.0%   ENYVAA---------SNENWSNAVHNLSGEWNTFTNSITA-----------------------
58 JADRTN010000001.1_ORF.648     79.2%  18.0%   ENYVAA---------SNENWSNAVHNLSGEWNTFTNSITA-----------------------
59 ALNJ01000086.1_ORF.55451      79.2%  20.7%   DSYVAA---------SNENWRNAVSDLSGEWNTFTNSITA-----------------------
60 CP020358.1_ORF.49881          79.2%  20.7%   ESYVAA---------SNENWSNAVHDLSGEWNTFTNSITA-----------------------
61 AKCF01000001.1_ORF.13999      79.2%  18.3%   IGSILADHLNSMMYEKSGIWSNFVYDAATNWGDVVSSLQK-----------------------
62 KI535631.1_ORF.67631          79.2%  18.3%   IGSILADHLNSMMYEKSGIWSNFVYDAATNWGDVVSSLQK-----------------------
63 ARVT01000001.1_ORF.83832      79.2%  18.3%   IGSILADHLNSMMYEKSGIWSNFVYDAATNWGDVVSSLQK-----------------------
64 JAKX01000001.1_ORF.726        79.2%  18.3%   IGSILADHLNSMMYEKSGIWSNFVYDAATNWGDVVSSLQK-----------------------
65 KK097709.1_ORF.46887          79.2%  18.3%   IGSILADHLNSMMYEKSGIWSNFVYDAATNWGDVVSSLQK-----------------------
66 BCZK01000002.1_ORF.16754      79.2%  18.3%   IGSILADHLNSMMYEKSGIWSNFVYDAATNWGDVVSSLQK-----------------------
67 DIEF01000003.1_ORF.23844      79.2%  18.3%   IGSILADHLNSMMYEKSGIWSNFVYDAATNWGDVVSSLQK-----------------------
68 KI535597.1_ORF.31267          79.2%  18.3%   IGSILADHLNSMMYEKSGIWSNFVYDAATNWGDVISSLQK-----------------------
69 JAKY01000041.1_ORF.44918      79.2%  18.3%   IGSILADHLNSMMYEKSGIWSNFVYDAATNWGDVISSLQK-----------------------
70 KQ235791.1_ORF.67988          79.2%  18.3%   IGSILADHLNSMMYEKSGIWSNFVYDAATNWGDVISSLQK-----------------------
71 JAND01000082.1_ORF.77829      76.7%  21.0%   TGAVIG-----------AILGRFVAGATTGAVTGASLDGVLFDQYECRDCEHTFD--------
72 JADRTN010000013.1_ORF.51264   76.7%  21.0%   TGAVIG-----------AILGRFVAGATTGAVTGASLDGVLFDQYECRDCEHTFD--------
73 AP022142.1_ORF.50706          76.7%  21.0%   TGAVIG-----------AILGRFVAGATAGAVTGASLDGVLFDQYECRDCEHTFD--------
74 AP022142.1_ORF.3785           76.7%  21.8%   AGAVIG-----------AVFGRFVAGATAGAVTGSALDGLVFDRYECGDCGHTFD--------
   consensus/100%                               .s.h.u...........t.ht.h..s..sths...t...........................
   consensus/90%                                .s.h.u...........t.hs.hh.sh.sths.h.t.h.t.......................
   consensus/80%                                .uhhhu...........t.hu.sltsh.sthshh.sphht.......................
   consensus/70%                                hGhlhu...........ulhu.ultsuhssWshhhsphht.......................
```

MView 1.63, Copyright © 1997-2018 Nigel P. Brown
