## Supplementary File S9 for "Semantic search using protein large language models detects class II microcins in bacterial genomes"

Reference sequence (1): KI973125.1_ORF.44026
Identities normalised by aligned length.
Colored by: identity

```
                                  cov     pid   1 [         .         .         .         :         .         .         . 80
 1 KI973125.1_ORF.44026         100.0%  100.0%  ------------------------MKAISFVEAKDIIGGAL--------------------NPFAG---LVK--------
 2 CP017183.1_ORF.209           100.0%  100.0%  ------------------------MKAISFVEAKDIIGGAL--------------------NPFAG---LVK--------
 3 WHPU01000006.1_ORF.66237     100.0%  100.0%  ------------------------MKAISFVEAKDIIGGAL--------------------NPFAG---LVK--------
 4 JADQTK010000001.1_ORF.107    100.0%  100.0%  ------------------------MKAISFVEAKDIIGGAL--------------------NPFAG---LVK--------
 5 FYBF01000109.1_ORF.61868     100.0%  100.0%  ------------------------MKAISFVEAKDIIGGAL--------------------NPFAG---LVK--------
 6 VKTL01000010.1_ORF.14872     100.0%   98.4%  ------------------------MKAISFVEAKDIIGGAL--------------------NPFAG---LVK--------
 7 VLMG01000003.1_ORF.51872     100.0%   98.4%  ------------------------MKAISFVEAKDIIGGAL--------------------NPFAG---LVK--------
 8 JADBPT010000026.1_ORF.47366  100.0%   98.4%  ------------------------MKAISFVEAKDIIGGAL--------------------NPFAG---LVK--------
 9 JADBPU010000017.1_ORF.45210  100.0%   98.4%  ------------------------MKAISFVEAKDIIGGAL--------------------NPFAG---LVK--------
10 JADQTI010000001.1_ORF.170    100.0%   98.4%  ------------------------MKAISFVEAKDIIGGAL--------------------NPFAG---LVK--------
11 JAFHGL010000133.1_ORF.10921  100.0%   98.4%  ------------------------MKAISFVEAKDIIGGAL--------------------NPFAG---LVK--------
12 KI973125.1_ORF.41812          95.3%   41.9%  ------------------------MQVIDFKKAQAIIGG-W--------------------DPFAA---AIQ--------
13 CP017183.1_ORF.23035          95.3%   41.9%  ------------------------MQVIDFKKAQAIIGG-W--------------------DPFAA---AIQ--------
14 VLMG01000003.1_ORF.51072      95.3%   41.9%  ------------------------MQVIDFKKAQAIIGG-W--------------------DPFAA---AIQ--------
15 WHPU01000006.1_ORF.65515      95.3%   41.9%  ------------------------MQVIDFKKAQAIIGG-W--------------------DPFAA---AIQ--------
16 JADBPT010000026.1_ORF.47188   95.3%   41.9%  ------------------------MQVIDFKKAQAIIGG-W--------------------DPFAA---AIQ--------
17 JADBPU010000017.1_ORF.44900   95.3%   41.9%  ------------------------MQVIDFKKAQAIIGG-W--------------------DPFAA---AIQ--------
18 JADQTI010000001.1_ORF.816     95.3%   41.9%  ------------------------MQVIDFKKAQAIIGG-W--------------------DPFAA---AIQ--------
19 JADQTK010000001.1_ORF.800     95.3%   41.9%  ------------------------MQVIDFKKAQAIIGG-W--------------------DPFAA---AIQ--------
20 FYBF01000109.1_ORF.61135      95.3%   41.9%  ------------------------MQVIDFKKAQAIIGG-W--------------------DPFAA---AIQ--------
21 VKTL01000010.1_ORF.15573      95.3%   41.9%  ------------------------MQVIDFKKAQAIIGG-W--------------------DPFAA---ALQ--------
22 JAFHGL010000133.1_ORF.10996   95.3%   41.9%  ------------------------MQVIDFKKAQAIIGG-W--------------------DPFAA---ALQ--------
23 JUZJ01000058.1_ORF.23041      98.4%   18.4%  M------------GGKMASCPNCKSTNVRERNVGKKTGGVI--------------------GGVGG---ALG--------
24 RSDS01000007.1_ORF.66833     100.0%   16.2%  -----------------MKCPDCGSTRVQRSDIGKKIGCGV--------------------GAVAG---GITGV---ISS
25 JAFBJM010000007.1_ORF.66835  100.0%   16.2%  -----------------MKCPDCGSTRVQRSDIGKKIGCGV--------------------GAVAG---GITGV---ISS
26 VLNN01000011.1_ORF.7068-23   100.0%   11.5%  ---------------MLDAWKVHEDDSLTPEQKKKQYATIT--------------------ARSAG---ASA--------
27 JWGM01000076.1_ORF.28259      81.2%   12.0%  M-------KKVLYGIFAISALAATSVYAAPVQVGE---------------------------AAGS---AAT--------
28 JABAIR010000007.1_ORF.60045   81.2%   12.0%  M-------KKVLYGIFAISALAATSVYAAPVQVGE---------------------------AAGS---AAT--------
29 WCIM01000006.1_ORF.6111       81.2%   10.9%  M-------KKVLYGIFAISALAATSVNAAPVQVGE---------------------------AAGS---AAT--------
30 CP056552.1_ORF.14054          81.2%   10.9%  M-------KKVLYGIFAISALAATSVNAAPVQVGE---------------------------AAGS---AAT--------
31 JABXRH010000001.1_ORF.43420   81.2%   10.9%  M-------KKVLYGIFAISALAATSVNAAPVQVGE---------------------------AAGS---AAT--------
32 FKEY01000011.1_ORF.49702      81.2%   10.9%  M-------KKVLYGIFAISALAATSVNAAPVQVGE---------------------------AAGS---AAT--------
33 CABGVW010000020.1_ORF.68950   81.2%   10.9%  M-------KKVLYGIFAISALAATSVNAAPVQVGE---------------------------AAGS---AAT--------
34 VLNO01000001.1_ORF.4161      100.0%   15.7%  M-------RTFFSGQMTRKADSTDSSHKGFVMKLMKTALIISTL-------IFSTSGMAIDKTAAG---AVA--------
35 RBXU01000015.1_ORF.63225     100.0%   13.9%  -MGEEIPEK----AIFTPESLSLVKGMAKAGRVVQVVGIVLTA---------YDHEQATEKSIKTG---SVKPISAEVIR
36 JABXRH010000001.1_ORF.32150   87.5%   11.4%  -------------------------------MGKSISKGF---------------------RSIAK---GLTGGA-----
37 AZUA01000011.1_ORF.61716      98.4%    9.5%  M---EKVYGYYGYTFCSLLQGTLIMRELNESELSSVSGAGM--------W-----------GSIGS---AIGGMFG----
38 RXPU01000017.1_ORF.23853      98.4%    9.5%  M---EKVYGYYGYTFCSLLQGTLIMRELNESELSNVSGAGM--------W-----------GSIGS---AIGGMFG----
39 CP035633.1_ORF.21695          98.4%    9.5%  M---EKVYGYYGYTFCSLLQGTLIMRELNESELSNVSGAGM--------W-----------GSIGS---AIGGMFG----
40 FJWF01000014.1_ORF.56787      98.4%    9.5%  M---EKVYGYYGYTFCSLLQGTLIMRELNESELSNVSGAGM--------W-----------GSIGS---AIGGMFG----
41 FJZP01000016.1_ORF.53830      98.4%    9.5%  M---EKVYGYYGYTFCSLLQGTLIMRELNESELSNVSGAGM--------W-----------GSIGS---AIGGMFG----
42 FKBI01000017.1_ORF.59533      98.4%    9.5%  M---EKVYGYYGYTFCSLLQGTLIMRELNESELSNVSGAGM--------W-----------GSIGS---AIGGMFG----
43 FKFV01000024.1_ORF.63613      98.4%    9.5%  M---EKVYGYYGYTFCSLLQGTLIMRELNESELSNVSGAGM--------W-----------GSIGS---AIGGMFG----
44 FKGO01000013.1_ORF.31834      98.4%    9.5%  M---EKVYGYYGYTFCSLLQGTLIMRELNESELSNVSGAGM--------W-----------GSIGS---AIGGMFG----
45 JAEKKB010000008.1_ORF.60800   98.4%    9.5%  M---EKVYGYYGYTFCSLLQGTLIMRELNECELSSVSGAGM--------W-----------GSIGS---AIGGMFG----
46 QFXN01000273.1_ORF.19861      98.4%    9.5%  M---EKVYGYYGYTFCSLLQGTLIMRELNESELSNVSGAGM--------W-----------GSIGS---AIGGMFG----
47 JDWG01000020.1_ORF.62376      98.4%    9.5%  M---EKVYGYYGYTFCSLLQGKLIMRELNENELSNVSGAGM--------W-----------GSIGS---AIGGMFG----
48 JDWH01000014.1_ORF.61217      98.4%    9.5%  M---EKVYGYYGYTFCSLLQGKLIMRELNENELSNVSGAGM--------W-----------GSIGS---AIGGMFG----
49 PZPP01000022.1_ORF.71288      98.4%    9.5%  M---EKVYGYYGYTFCSLLQGKLIMRELNENELSNVSGAGM--------W-----------GSIGS---AIGGMFG----
50 RSDS01000013.1_ORF.6611       98.4%    9.5%  M---EKVYGYYGYTFCSLLQGKLIMRELNENELSNVSGAGM--------W-----------GSIGS---AIGGMFG----
51 JAFBJM010000013.1_ORF.27400   98.4%    9.5%  M---EKVYGYYGYTFCSLLQGKLIMRELNENELSNVSGAGM--------W-----------GSIGS---AIGGMFG----
52 JUZJ01000011.1_ORF.72885      98.4%    9.5%  M---EKVYGYYGYTFCSLLQGKFIMRELNENELSNVSGAGM--------W-----------GSIGS---AIGGMFG----
53 CP056394.1_ORF.77065-5        98.4%    5.9%  MIATMTPAGMLAGAVLVGDALNTVSREAARQFLNEPASEGI---------------------LADG---AMS-----VAE
54 CABGVW010000021.1_ORF.71043-6 98.4%    6.0%  MMSSMTPVGVVAGAVIVVNGFNTISREAAHNLLGDKQTEGI---------------------FADG---SME-----FAE
55 AZUA01000004.1_ORF.43055-25   98.4%    7.6%  M------------SIYQRYLANQSPVPLNLVAPEDIVDMGV--------------------DSAGN---FVCG-------
56 JAEKKB010000004.1_ORF.48579-25 98.4%   7.6%  M------------SIYQRYLANQSPVPLNLVAPEDIVDMGV--------------------DSAGN---FVCG-------
57 FKEY01000003.1_ORF.16130-25   98.4%    7.6%  M------------SIYQRYLANQSPVPLNLVAPEDIVDMGV--------------------DGSGN---FVCG-------
58 JACRRJ010000003.1_ORF.79249-6 87.5%    9.2%  M-------------------------------LPDNEAQLL--------------------STSDNKIISFSGSHM--SF
59 JACRRJ010000004.1_ORF.79880-6 87.5%    9.2%  M-------------------------------LPDNEAQLL--------------------STSDNKIISFSGSHM--SF
60 FJZP01000035.1_ORF.74011-3   100.0%    6.7%  M----------STVMVLSAVLFGAGRKGLNLLKDKDTGNIYEIRGSKAYRLSDEEAVRYQTSMSGKGIALAEFSS--SQ
61 CABGKT010000038.1_ORF.60701   76.6%    6.3%  M----------------------------SKLLRELTDKEI-------------------QSSLSN--------------
62 WCIM01000016.1_ORF.27377      78.1%   17.2%  M--------------------------------KRFISVALLA------------------ALLAG--CAHDSPCVPVYD
63 JAEKKB010000002.1_ORF.39405   78.1%   17.2%  M--------------------------------KRFISVALLA------------------ALLAG--CAHDSPCVPVYD
   consensus/100%                               ................................................................st..............
   consensus/90%                                ...........................ht...htt...........................shus...hht........
   consensus/80%                                ...........................hs..thtph.sh.h....................sshuu...hht........
   consensus/70%                                .........................p.ls..phpsh.us.h....................sshuu...ult........

                                  cov     pid  81         .         1         .         .         .         .         :         . 160
 1 KI973125.1_ORF.44026         100.0%  100.0%  --GAQLGYDT-GASIMGMVGGV------VGGVLGG-------------------------AMGFLGA-LVGSYN------
 2 CP017183.1_ORF.209           100.0%  100.0%  --GAQLGYDT-GASIMGMVGGV------VGGVLGG-------------------------AMGFLGA-LVGSYN------
 3 WHPU01000006.1_ORF.66237     100.0%  100.0%  --GAQLGYDT-GASIMGMVGGV------VGGVLGG-------------------------AMGFLGA-LVGSYN------
 4 JADQTK010000001.1_ORF.107    100.0%  100.0%  --GAQLGYDT-GASIMGMVGGV------VGGVLGG-------------------------AMGFLGA-LVGSYN------
 5 FYBF01000109.1_ORF.61868     100.0%  100.0%  --GAQLGYDT-GASIMGMVGGV------VGGVLGG-------------------------AMGFLGA-LVGSYN------
 6 VKTL01000010.1_ORF.14872     100.0%   98.4%  --GAQLGYDT-GASIMGMVGGV------VGGILGG-------------------------AMGFLGA-LVGSYN------
 7 VLMG01000003.1_ORF.51872     100.0%   98.4%  --GAQLGYDT-GASIMGMVGGV------VGGILGG-------------------------AMGFLGA-LVGSYN------
 8 JADBPT010000026.1_ORF.47366  100.0%   98.4%  --GAQLGYDT-GASIMGMVGGV------VGGILGG-------------------------AMGFLGA-LVGSYN------
 9 JADBPU010000017.1_ORF.45210  100.0%   98.4%  --GAQLGYDT-GASIMGMVGGV------VGGILGG-------------------------AMGFLGA-LVGSYN------
10 JADQTI010000001.1_ORF.170    100.0%   98.4%  --GAQLGYDT-GASIMGMVGGV------VGGILGG-------------------------AMGFLGA-LVGSYN------
11 JAFHGL010000133.1_ORF.10921  100.0%   98.4%  --GAQLGYDT-GASIMGMVGGV------VGGILGG-------------------------AMGFLGA-LVGSYN------
12 KI973125.1_ORF.41812          95.3%   41.9%  --GASAGYNA-GSQMLGALGGT------IGGVFGL-------------------------VGGFIGG-FFSA--------
13 CP017183.1_ORF.23035          95.3%   41.9%  --GASAGYNA-GSQMLGALGGT------IGGVFGL-------------------------VGGFIGG-FFSA--------
14 VLMG01000003.1_ORF.51072      95.3%   41.9%  --GASAGYNA-GSQMLGALGGT------IGGVFGL-------------------------VGGFIGG-FFSA--------
15 WHPU01000006.1_ORF.65515      95.3%   41.9%  --GASAGYNA-GSQMLGALGGT------IGGVFGL-------------------------VGGFIGG-FFSA--------
16 JADBPT010000026.1_ORF.47188   95.3%   41.9%  --GASAGYNA-GSQMLGALGGT------IGGVFGL-------------------------VGGFIGG-FFSA--------
17 JADBPU010000017.1_ORF.44900   95.3%   41.9%  --GASAGYNA-GSQMLGALGGT------IGGVFGL-------------------------VGGFIGG-FFSA--------
18 JADQTI010000001.1_ORF.816     95.3%   41.9%  --GASAGYNA-GSQMLGALGGT------IGGVFGL-------------------------VGGFIGG-FFSA--------
```

```
19 JADQTK010000001.1_ORF.800      95.3%  41.9%   --GASAGYNA-GSQMLGALGGT------IGGVFGL-------------------------VGGFIGG-FFSA--------
20 FYBF01000109.1_ORF.61135       95.3%  41.9%   --GASAGYNA-GSQMLGALGGT------IGGVFGL-------------------------VGGFIGG-FFSA--------
21 VKTL01000010.1_ORF.15573       95.3%  41.9%   --GASAGYNA-GSQMLGALGGT------IGGVFGL-------------------------VGGFIGG-FFSA--------
22 JAFHGL010000133.1_ORF.10996    95.3%  41.9%   --GASAGYNA-GSQMLGALGGT------IGGVFGL-------------------------VGGFIGG-FFSA--------
23 JUZJ01000058.1_ORF.23041       98.4%  18.4%   --GASAGAAV-GS-IVPGIGTA------VGAVVGGLL----------------GRFAAASAGAAAGA-VTGSALDGIVFD
24 RSDS01000007.1_ORF.66833      100.0%  16.2%   REGVEAGVAI-GSAILPGIGTI------AGGVLGLLI--------------------GVGSGAVIGG-LAGEAVDDHLFN
25 JAFBJM010000007.1_ORF.66835   100.0%  16.2%   REGVEAGVAI-GSAILPGIGTI------AGGVLGLLI--------------------GVGSGAVIGG-LAGEAVDDHLFN
26 VLNN01000011.1_ORF.7068-23    100.0%  11.5%   --GGEAGATM-GTMVFPGVGTI------LGGIIGS-------------------------IGGYYGGDLIGDKLPDSLFT
27 JWGM01000076.1_ORF.28259       81.2%  12.0%   --SASAGSST-AA-SASTVSSA------VGV-------------------------ALAATGGGDGS-NTGTTTTTTTST
28 JABAIR010000007.1_ORF.60045    81.2%  12.0%   --SASAGSST-AA-SASTVSSA------VGV-------------------------ALAATGGGDGS-NTGTTTTTTTST
29 WCIM01000006.1_ORF.6111        81.2%  10.9%   --SASAGIST-AS-SASTVSSA------VGV-------------------------ALAATGGGDGS-NTGTTTTTTTST
30 CP056552.1_ORF.14054           81.2%  10.9%   --SASAGIST-AS-SASTVSSA------VGV-------------------------ALAATGGGDGS-NTGTTTTTTTST
31 JABXRH010000001.1_ORF.43420    81.2%  10.9%   --SASAGIST-AS-SASTVSSA------VGV-------------------------ALAATGGGDGS-NTGTTTTTTTST
32 FKEY01000011.1_ORF.49702       81.2%  10.9%   --SASAGIST-AS-SASTVSSA------VGV-------------------------ALAATGGGDGS-NTGTTTTTTTST
33 CABGVW010000020.1_ORF.68950    81.2%  10.9%   --SASAGIST-AS-SASTVSSA------VGV-------------------------ALAATGGGDGS-NTGTTTTTTTST
34 VLNO01000001.1_ORF.4161       100.0%  15.7%   --GAAIGAAT-GKDLKSTVGGA-----VVGAGTGAMFKN-------------GDKGKAARKGGAVGA-VVGAGAAAVTGN
35 RBXU01000015.1_ORF.63225      100.0%  13.9%   QAGGWGGFVA-GAKIGTVAGAA------VGVETGP------------------GAVITGLIGGIIFG-CAGYYGADWIAD
36 JABXRH010000001.1_ORF.32150    87.5%  11.4%   --GTQGGASV-PPPVAPPVAPPDDAQQAVVDTQGQETEG-------------GKTMSRGKSGLNISR-TSGGGLSL----
37 AZUA01000011.1_ORF.61716       98.4%   9.5%   --GGSAGCN--GSYSHTSTTNKDGSHSESTSFSGG---------------------CSVNVGGKSGGSSKGNGGSKSGGG
38 RXPU01000017.1_ORF.23853       98.4%   9.5%   --GGSAGCN--GSYSHTSTTNKDGSHSESTSFSGG---------------------CSVNVGGKSGGSSKGNGGSKSGGG
39 CP035633.1_ORF.21695           98.4%   9.5%   --GGSAGCN--GSYSHTSTTNKDGSHSESTSFSGG---------------------CSVNVGGKSGGSSKGNGGSKSGGG
40 FJWF01000014.1_ORF.56787       98.4%   9.5%   --GGSAGCN--GSYSHTSTTNKDGSHSESTSFSGG---------------------CSVNVGGKSGGSSKGNGGSKSGGG
41 FJZP01000016.1_ORF.53830       98.4%   9.5%   --GGSAGCN--GSYSHTSTTNKDGSHSESTSFSGG---------------------CSVNVGGKSGGSSKGNGGSKSGGG
42 FKBI01000017.1_ORF.59533       98.4%   9.5%   --GGSAGCN--GSYSHTSTTNKDGSHSESTSFSGG---------------------CSVNVGGKSGGSSKGNGGSKSGGG
43 FKFV01000024.1_ORF.63613       98.4%   9.5%   --GGSAGCN--GSYSHTSTTNKDGSHSESTSFSGG---------------------CSVNVGGKSGGSSKGNGGSKSGGG
44 FKGO01000013.1_ORF.31834       98.4%   9.5%   --GGSAGCN--GSYSHTSTTNKDGSHSESTSFSGG---------------------CSVNVGGKSGGSSKGNGGSKSGGG
45 JAEKKB010000008.1_ORF.60800    98.4%   9.5%   --GGSAGCN--GSYSHTSTTNKDGSHSESTSFSGG---------------------CSVNVGGKSGGSSKGNGGSKSGGG
46 QFXN01000273.1_ORF.19861       98.4%   9.5%   --GGSAGCN--GSYSHTSTTNKDGSHSESTSFSGG---------------------CSVNVGGKSGGSSKGNGGSKSGGG
47 JDWG01000020.1_ORF.62376       98.4%   9.5%   --GGSAGCN--GSYSHTSTTNKDGSHSESTSFSGG---------------------CSVNVGGKSGGSSKGNGGSKNGGG
48 JDWH01000014.1_ORF.61217       98.4%   9.5%   --GGSAGCN--GSYSHTSTTNKDGSHSESTSFSGG---------------------CSVNVGGKSGGSSKGNGGSKNGGG
49 PZPP01000022.1_ORF.71288       98.4%   9.5%   --GGSAGCN--GSYSHTSTTNKDGSHSESTSFSGG---------------------CSVNVGGKSGGSSKGNGGSKNGGG
50 RSDS01000013.1_ORF.6611        98.4%   9.5%   --GGSAGCN--GSYSHTSTTNKDGSHSESTSFSGG---------------------CSVNVGGKSGGSSKGNGGSKNGGG
51 JAFBJM010000013.1_ORF.27400    98.4%   9.5%   --GGSAGCN--GSYSHTSTTNKDGSHSESTSFSGG---------------------CSVNVGGKSGGSSKGNGGSKNGGG
52 JUZJ01000011.1_ORF.72885       98.4%   9.5%   --GGSAGCN--GSYSHTSTTNKDGSHSESTSFSGG---------------------CSVNVGGKSGGSSKGNGGSKNGGG
53 CP056394.1_ORF.77065-5         98.4%   5.9%   FMGFSRDVGL-GIFNATTLAAS------VYGVFGLTGKAEAWRLFRYLPGDYYRKVETMSPGKLTMK-VLGWGSVQRWYW
54 CABGVW010000021.1_ORF.71043-6  98.4%   6.0%   FMGFSRQQGL-GTYKAVTLFSE------VYGAYGLRLKPEAQRLWYWTRPDFFRKVSGTPRPAMALK-IAGWGVKAKVAL
55 AZUA01000004.1_ORF.43055-25    98.4%   7.6%   --GGMRTLPV-WTANPDSLCQQSQPEQPTGNPFDQSSQPQQPQ--QQQPQQQNEKKDSDGVAGWIKD-MFGGN-------
56 JAEKKB010000004.1_ORF.48579-25 98.4%   7.6%   --GGMRTLPV-WTANPDSLCQQSQPEQPTGNPFDQSSQPQQPQ--QQQPQQQNEKKDSDGVAGWIKD-MFGGN-------
57 FKEY01000003.1_ORF.16130-25    98.4%   7.6%   --GGMRTLPV-WTTNPDSLCQQSQPEQPTGNPFEQSSQPQQQP--QQQPQQQNEKKDSDGVAGWIKD-MFGGN-------
58 JACRRJ010000003.1_ORF.79249-6  87.5%   9.2%   QDGGITNIPL-NTSFNDSFSNG----ITVNPASGMPM-----------------------VGGISGLDIHGNSWGTNFNE
59 JACRRJ010000004.1_ORF.79880-6  87.5%   9.2%   QDGGITNIPL-NTSFNDSFSNG----ITVNPASGMPM-----------------------VGGISGLDIHGNSWGTNFNE
60 FJZP01000035.1_ORF.74011-3    100.0%   6.7%   YSGLDTNSSVLPINFGPSHTDI-NQGIAINPSSGMPM-----------------------VGGISGLDIHGNSWGTNFNE
61 CABGKT010000038.1_ORF.60701    76.6%   6.3%   --GGSTCSMY-KALLPGHIAAC-------------------------------------AASAYFYG-SSGCTAMRYVSE
62 WCIM01000016.1_ORF.27377       78.1%  17.2%   DQGRLVHTNT------CMKGTTQDNWETAGAIAGG----------------------AAAVAGLTLG-IVALTK------
63 JAEKKB010000002.1_ORF.39405    78.1%  17.2%   DQGRLVHTNT------CMKGTTQDNWETAGAIAGG----------------------AAAVAGLTLG-IVALTK------
   consensus/100%                                ..uh.ht.........s.h...........................................h.......u.........
   consensus/90%                                 ..us.hshs..ss...s.hssh.......ss.............................shuh.ts..hut........
   consensus/80%                                 ..GuphGhs..us..hs.hssh.......ss..G..........................suGh.Gu..hus........
   consensus/70%                                 ..GupsGhsh.Gu..hsshsst......ssuh.Gh.........................sGGh.Gu..hGsh.......

                                    cov    pid 161          .          .          .          2         ] 210
 1 KI973125.1_ORF.44026          100.0% 100.0%   --------------------------------------------------
 2 CP017183.1_ORF.209            100.0% 100.0%   --------------------------------------------------
 3 WHPU01000006.1_ORF.66237      100.0% 100.0%   --------------------------------------------------
 4 JADQTK010000001.1_ORF.107     100.0% 100.0%   --------------------------------------------------
 5 FYBF01000109.1_ORF.61868      100.0% 100.0%   --------------------------------------------------
 6 VKTL01000010.1_ORF.14872      100.0%  98.4%   --------------------------------------------------
 7 VLMG01000003.1_ORF.51872      100.0%  98.4%   --------------------------------------------------
 8 JADBPT010000026.1_ORF.47366   100.0%  98.4%   --------------------------------------------------
 9 JADBPU010000017.1_ORF.45210   100.0%  98.4%   --------------------------------------------------
10 JADQTI010000001.1_ORF.170     100.0%  98.4%   --------------------------------------------------
11 JAFHGL010000133.1_ORF.10921   100.0%  98.4%   --------------------------------------------------
12 KI973125.1_ORF.41812           95.3%  41.9%   --------------------------------------------------
13 CP017183.1_ORF.23035           95.3%  41.9%   --------------------------------------------------
14 VLMG01000003.1_ORF.51072       95.3%  41.9%   --------------------------------------------------
15 WHPU01000006.1_ORF.65515       95.3%  41.9%   --------------------------------------------------
16 JADBPT010000026.1_ORF.47188    95.3%  41.9%   --------------------------------------------------
17 JADBPU010000017.1_ORF.44900    95.3%  41.9%   --------------------------------------------------
18 JADQTI010000001.1_ORF.816      95.3%  41.9%   --------------------------------------------------
19 JADQTK010000001.1_ORF.800      95.3%  41.9%   --------------------------------------------------
20 FYBF01000109.1_ORF.61135       95.3%  41.9%   --------------------------------------------------
21 VKTL01000010.1_ORF.15573       95.3%  41.9%   --------------------------------------------------
22 JAFHGL010000133.1_ORF.10996    95.3%  41.9%   --------------------------------------------------
23 JUZJ01000058.1_ORF.23041       98.4%  18.4%   KYICRD----------------------------CEHTFD----------
24 RSDS01000007.1_ORF.66833      100.0%  16.2%   TYCCDV----------------------------CGGQFEGRIS------
25 JAFBJM010000007.1_ORF.66835   100.0%  16.2%   TYCCDV----------------------------CGGQFEGRIS------
26 VLNN01000011.1_ORF.7068-23    100.0%  11.5%   RSVSNKKQDTITPMDVLGLNPQRLDVHLYVDGQEIHAALDDRTNRDALRN
27 JWGM01000076.1_ORF.28259       81.2%  12.0%   Q-------------------------------------------------
28 JABAIR010000007.1_ORF.60045    81.2%  12.0%   Q-------------------------------------------------
29 WCIM01000006.1_ORF.6111        81.2%  10.9%   Q-------------------------------------------------
30 CP056552.1_ORF.14054           81.2%  10.9%   Q-------------------------------------------------
31 JABXRH010000001.1_ORF.43420    81.2%  10.9%   Q-------------------------------------------------
32 FKEY01000011.1_ORF.49702       81.2%  10.9%   Q-------------------------------------------------
33 CABGVW010000020.1_ORF.68950    81.2%  10.9%   Q-------------------------------------------------
34 VLNO01000001.1_ORF.4161       100.0%  15.7%   SVLKGAAVGA------------------------GSGALIGEAT-----H
35 RBXU01000015.1_ORF.63225      100.0%  13.9%   YIDEN---------------------------------------------
36 JABXRH010000001.1_ORF.32150    87.5%  11.4%   --------------------------------------------------
37 AZUA01000011.1_ORF.61716       98.4%   9.5%   RQGNDHG-----------------------------------------RH
38 RXPU01000017.1_ORF.23853       98.4%   9.5%   RQGNDHG-----------------------------------------RH
39 CP035633.1_ORF.21695           98.4%   9.5%   RQGNDHG-----------------------------------------RH
40 FJWF01000014.1_ORF.56787       98.4%   9.5%   RQGNDHG-----------------------------------------RH
41 FJZP01000016.1_ORF.53830       98.4%   9.5%   RQGNDHG-----------------------------------------RH
42 FKBI01000017.1_ORF.59533       98.4%   9.5%   RQGNDHG-----------------------------------------RH
43 FKFV01000024.1_ORF.63613       98.4%   9.5%   RQGNDHG-----------------------------------------RH
```

```
44 FKGO01000013.1_ORF.31834        98.4%   9.5%     RQGNDHG-----------------------------------------RH
45 JAEKKB010000008.1_ORF.60800     98.4%   9.5%     RQGNDHG-----------------------------------------RH
46 QFXN01000273.1_ORF.19861        98.4%   9.5%     HQGNDHG-----------------------------------------RH
47 JDWG01000020.1_ORF.62376        98.4%   9.5%     RQGNDHG-----------------------------------------RH
48 JDWH01000014.1_ORF.61217        98.4%   9.5%     RQGNDHG-----------------------------------------RH
49 PZPP01000022.1_ORF.71288        98.4%   9.5%     RQGNDHG-----------------------------------------RH
50 RSDS01000013.1_ORF.6611         98.4%   9.5%     RQGNDHG-----------------------------------------RH
51 JAFBJM010000013.1_ORF.27400     98.4%   9.5%     RQGNDHG-----------------------------------------RH
52 JUZJ01000011.1_ORF.72885        98.4%   9.5%     RQGNDHG-----------------------------------------RH
53 CP056394.1_ORF.77065-5          98.4%   5.9%     NSLPVM----------------------------TGIADM----------
54 CABGVW010000021.1_ORF.71043-6   98.4%   6.0%     DLLTIE----------------------------PSGK------------
55 AZUA01000004.1_ORF.43055-25     98.4%   7.6%     --------------------------------------------------
56 JAEKKB010000004.1_ORF.48579-25  98.4%   7.6%     --------------------------------------------------
57 FKEY01000003.1_ORF.16130-25     98.4%   7.6%     --------------------------------------------------
58 JACRRJ010000003.1_ORF.79249-6   87.5%   9.2%     PSNTYD----------------------------PNRGY-----------
59 JACRRJ010000004.1_ORF.79880-6   87.5%   9.2%     PSNTYD----------------------------PNRGY-----------
60 FJZP01000035.1_ORF.74011-3     100.0%   6.7%     PSNTYD----------------------------PNRGY-----------
61 CABGKT010000038.1_ORF.60701     76.6%   6.3%     YIRKNC--------------------------------------------
62 WCIM01000016.1_ORF.27377        78.1%  17.2%     --------------------------------------------------
63 JAEKKB010000002.1_ORF.39405     78.1%  17.2%     --------------------------------------------------
   consensus/100%                                   ..................................................
   consensus/90%                                    ..................................................
   consensus/80%                                    ..................................................
   consensus/70%                                    ..................................................
```

MView 1.63, Copyright © 1997-2018 Nigel P. Brown
